# Cracking the Capsid Code: A Computationally-Feasible Approach for Investigating Virus-Excipient Interactions in Biologics Design

**DOI:** 10.1101/2025.09.04.674344

**Authors:** Jonathan W. P. Zajac, Idris Tohidian, Praveen Muralikrishnan, Sarah L. Perry, Caryn L. Heldt, Sapna Sarupria

## Abstract

The efficacy and equitable distribution of viral biologics, including vaccines and virus-like particles, is hindered due to their inherently low shelf life. To increase the longevity of such products, formulations are typically developed with small molecule additives known as excipients. Finding the correct excipients for a biological formulation is a costly and time-consuming process due to the large excipient design space and unknown mechanisms underlying excipient-virus interactions. Molecular dynamics simulations are, in theory, well-equipped to efficiently investigate these mechanisms. However, the massive size of fully assembled viral capsids, the protein shell that encapsulates the viral genome, demands computational resources well beyond the requirements of conventional simulations. There exists a need for a novel method that enables high-throughput investigations of virus-excipient interactions at the molecular level and at atomistic resolution. Here, we introduce **CapSACIN** — a computational framework for **Cap**sid **S**urface **A**bstraction and **C**omputationally-**I**nduced **N**anofragmentation. We demonstrate the applicability of this workflow to a model non-enveloped virus, porcine parvovirus (PPV). Through simulations of PPV surface models, we observe that the 2-fold axis of symmetry is significantly weaker at the molecular level than the 3- or 5-fold axes of symmetry. Further, we present results demonstrating excellent agreement with experimentally determined excipient effects on PPV thermal stability.

## 1 Introduction

Viral-based biologics are designed to either trigger an immune response ^1–7^ or to encapsulate cargo through the use of virus-like particles (VLPs).^8–13^ The primary ingredient in such formulations is the virus capsid; a protein shell that protects the infectious viral genome. VLPs are biomimetic materials that retain many inherent properties of virus capsids, including size, biocompatibility, stability, and cell entry behavior. ^14,15^ Repurposed to encapsulate both natural and synthetic cargo,^8,15^ VLPs have been used for intracellular delivery,^16^ imaging,^17^ and biocatalytic nanocompartments.^18–20^

Most viral therapeutics have recommended storage temperatures of 2–8 ^◦^C and have inherently low shelf lives.^21–23^ Potency loss at elevated temperatures is driven by the denaturation of capsid proteins, ^24,25^ disassembly of the capsid shell, ^26,27^ or aggregation of viral particles.^28,29^ Because of this temperature constraint, the development and distribution of viral biologics is confined within a “cold chain”. ^30–34^ The cold chain preserves viral vectors at refrigerated temperatures, but commonly fails due to improper training and equipment malfunction.^35–37^ Breakdowns in the cold chain give rise to loss of product integrity and lead directly to significant economical and societal burden. Hence, strategies that ease or eliminate the burden of the cold chain are paramount for widespread biologics distribution and reducing waste.

Inspired by the phenomenon of intracellular osmolyte production,^38–43^ excipient incorporation into formulations is perhaps the most common strategy for biologics stabilization. During formulation design, excipient selection is high-throughput and empirical for two key reasons.^44–47^ First, the excipient design space is extremely large. Most biological formulations include 5 or more unique excipients^48^ from a pool of ∼1000 possible excipients.^49^ Second, no single excipient can universally stabilize all virus capsids; excipient mechanisms, in general, are poorly understood. This gives rise to a trial-and-error process, bottlenecking formulation development.

Excipient-protein interactions (e.g., electrostatic, hydrogen-bonding, or van der Waals interactions) are thought to underlie the stabilization of viruses and VLPs,^50–52^ as these interactions are closely linked to changes in protein flexibility, structure, and stability.^45,53–60^ Molecular dynamics (MD) simulations offer a powerful means of investigating virus-excipient interactions by capturing their spatial and temporal behavior in atomistic detail. Simulations are also less resource-intensive than wet-lab experiments, reducing reliance on traditional, time-consuming approaches to excipient selection.

Atomistic simulations of individual proteins are well within the reach of modern computational power, though virus simulations are more computationally demanding. Virus capsids are comprised of 60 or more repeating protein subunits, with even the smallest capsids reaching 10-30 nm in diameter and 200-700 amino acid residues per monomer.^61^ While simulations of isolated capsid proteins are computationally tractable, they lack key features of assembled capsids such as curvature, ^62,63^ asymmetric dynamics,^64,65^ protein-protein interactions,^66^ and inter-protein binding sites.^67^ Hence, an accurate representation of viral capsids requires the full molecular context of the virus. To achieve meaningful atomistic simulations of fully assembled viral capsids, substantial dedicated hardware and computational time is required (Table 1). Brute-force computational power is still not sufficient for more complex viruses, such as enveloped viruses, which demand multiscale resolution or coarse-grained representations to achieve meaningful performance. ^68–70^ While these methods can provide significant computational speed-ups, ^71,72^ accurate mechanistic insights into virus stability rely on capturing the atomistic details of excipient–water–capsid interactions.

**Table 1:** Simulation performance and computational cost associated with recent atomistic virus simulations.

| Virus | Size<br>[atoms] | Resources<br>[Cores / GPUs] | Performance<br>[ns/day] | Year | Reference |
| --- | --- | --- | --- | --- | --- |
| STMV | 1.2M | 2,016 / 0 | 30.0 | 2012 | Larsson *et al.*<sup>73</sup> |
| PV | 2.75M | 2048 / 0 | 1.27 | 2012 | Roberts *et al.*<sup>74</sup> |
| AMCV | 4M | 128 / 0 | 0.3 | 2014 | Arcangeli *et al.*<sup>75</sup> |
| HIV-1 | 64M | 62,080 / 3,880 | Not Listed | 2017 | Perilla and Schulten<sup>76</sup> |
| EV-1 | 3.2M | 2,048 / 0 | 11.0 | 2017 | Pohjolainen *et al.*<sup>77</sup> |
| IAV | 160M | 114,688 / 0 | 4.42 | 2020 | Durrant *et al.*<sup>68</sup> |
| SARS-CoV-2 | 305M | 180,224 / 24,576 | 68.36 | 2020 | Casalino *et al.*<sup>78</sup> |
| FMDV Subunit | 350K | 5,360 / 24 | 24.9 | 2021 | Li *et al.*<sup>79</sup> |
| H1N1 (CG) | 160M | 28,672 / 0 | 13.83 | 2022 | Casalino *et al.*<sup>69</sup> |
| SARS-CoV-2 (CG) | 31M | 3,920 / 0 | 10.0 | 2023 | Wang *et al.*<sup>72</sup> |
| SARS-CoV-2 (RA) | 1B | 180,224 / 24,576 | 34.1 | 2023 | Dommer *et al.*<sup>80</sup> |
| FHV | 5M | 400 / 20 | 15.5 | 2024 | Shrivastav *et al.*<sup>81</sup> |

Abbreviations: STMV – satellite tobacco mosaic virus; PV – poliovirus; AMCV – artichoke mottled crinkle virus; HIV-1 - human immunodeficiency virus type 1; EV-1 – echovirus type 1; IAV – influenza virus type A; SARS-CoV-2 – severe acute respiratory syndrome coronavirus 2; FMDV – foot-and-mouth disease virus; FHV – flock house virus; CG – coarse-grained; RA – respiratory aerosol.

There exists an unmet need in biologics design for high-throughput—yet high-detail—investigations of virus-excipient interactions. We address this need by developing a method that simplifies fully assembled capsid structures into models that (i) are computationally feasible, (ii) accurately reproduce capsid structure and dynamics, (iii) retain atomistic resolution, and (iv) can efficiently evaluate excipient stabilizing effects. To this end, we introduce a novel framework: **CapSACIN** — a computational framework for **Cap**sid **S**urface **A**bstraction and **C**omputationally-**I**nduced **N**anofragmentation. This method enables high-throughput identification of virus–excipient interactions at atomistic detail. CapSACIN leverages the symmetry of icosahedral capsids to construct surface models of viruses focused on a region of interest (ROI) embedded in peripheral proteins representing the molecular context of an assembled capsid. We further utilize a computational nanofragmentation approach, inspired by Ghaemi *et al.*,^82^ to assess the interfacial stability of monomer-monomer interfaces spanning the capsid surface. Using the non-enveloped porcine parvovirus (PPV) as a model system, we demonstrate that CapSACIN can accurately predict excipients as stabilizing or destabilizing, as validated by wet-lab experiments. In addition to predictive power, these simulations reveal mechanistic insights into excipient effects on weak interprotein interfaces. By bridging atomistic resolution with high-throughput capability, CapSACIN offers a powerful, information-driven strategy for accelerating formulation design and enhancing the development of stable, globally accessible viral biologics.

## 2 Methods

### 2.1 CapSACIN Workflow

CapSACIN is an end-to-end workflow capable of preparing input for and running capsid-excipient simulations (Fig. 1), composed of six steps: *Step 1: Symmetry-Based Alignment* (Fig. 1a), which involves aligning the ROI perpendicular to the *z*-axis and embedding it within the appropriate context; *Step 2: Surface Abstraction* (Fig. 1b), for abstracting the ROI and peripheral, context-providing proteins in preparation of simulations; *Step 3: Position Restraint Generation* (Fig. 1c), where an implicit wall is placed and position restraints are applied based on distance from the wall (to preserve capsid structural integrity and prevent excipient diffusion into the capsid interior); *Step 4: Restrained Simulations* (Fig. 1d), where applied restraints allow the solvent to equilibrate while preserving capsid structure; *Step 5: Unrestrained Simulations* (Fig. 1e), which allows the capsid surface to equilibrate in response to the solvent; and *Step 6: Nanofragmentation Simulations* (Fig. 1f), which enables prediction of stabilizing/destabilizing excipient effects on capsid protein interfaces.

**Figure 1:**
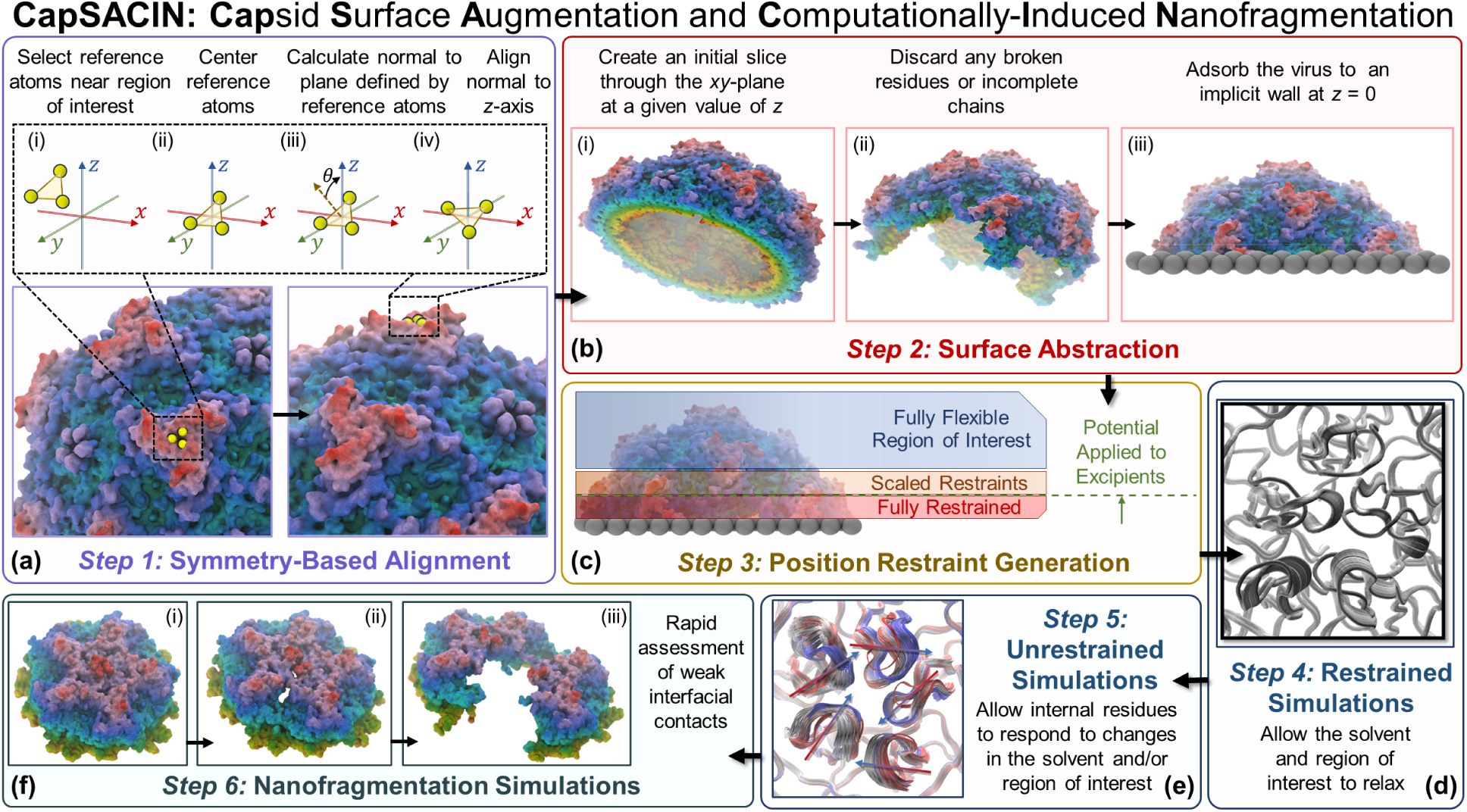
CapSACIN workflow. (a) *Step 1:* Symmetry-Based Alignment. (b) *Step 2:* Surface Abstraction. (c) *Step 3:* Position Restraint Generation. (d) *Step 4:* Restrained Simulations. (e) *Step 5:* Unrestrained Simulations. (f) *Step 6:* Nanofragmentation Simulations. All renders of molecular systems were completed in VMD. ^83^

The CapSACIN workflow depends only on an input PDB file, the selection of at least one reference atom within the ROI, and the target axis of symmetry. Details of each step in the CapSACIN workflow are provided below. Validations of a proof-of-concept surface model, including equilibration, structural integrity, and dynamics, were compared against the fully assembled PPV capsid.

#### 2.1.1 *Step 1:* Symmetry-Based Alignment

The first stage of the structure construction module involves a symmetry-based alignment of an input capsid structure (Fig. 1a). This stage requires only an input PDB file, a selected atom as a reference, and a desired symmetry. Capsid Cartesian coordinates are transformed such that the center-of-geometry of reference atoms is at the origin. Following translation, the normal to the plane connecting the three reference atoms is rotated such that it becomes aligned with the *z*-axis. For further details, see SI Section 1.

#### 2.1.2 *Step 2:* Surface Abstraction

Following alignment, the capsid structure is converted into a surface model (Fig. 1b). In this stage, the capsid is shifted such that its center-of-geometry is at the origin. A crude initial slice is made through the *xy*-plane at a height defined by a weight parameter *ω* ∈ [0, 1], where *ω* = 0 retains the full capsid and *ω* = 1 removes all capsid atoms. Atoms with *r_z_ < ωm_z_* are deleted, where *r_z_* is the *z*-coordinate of an atom and *m_z_* is the maximum *z*-coordinate of the capsid. For further details, see SI Section 1. This operation leaves dangling bonds, broken residues, and incomplete capsid proteins, so a clean-up step follows. Residues are removed if the number of heavy atoms is less than the expected count for a given residue type. Similarly, CapSACIN removes monomers if they contain fewer residues than expected from the input structure. The remaining capsid is translated upward such that its minimum *z*-coordinate is 0.

Capsid alignment and surface abstraction can, in theory, be extended to any icosahedral (or similarly structured), non-enveloped virus with an experimentally-resolved structure. In demonstrating our approach, we focus on the PPV capsid (PDB: 1K3V),^84^ though we have tested this stage of the CapSACIN workflow on a diverse set of capsid architectures (Fig. S1). Heuristically, we selected *ω* = 0.7 for PPV, as this provided a 2:1 ratio of peripheral to ROI proteins at the 5-, 3-, and 2-fold axes of symmetry. This guarantees three levels of capsid proteins: (i) terminal peripheral proteins that do not interact with the ROI; (ii) peripheral proteins sandwiched between ROI proteins and terminal peripheral proteins; and (iii) ROI proteins that interact with non-terminal peripheral proteins. In Section 2.1.3, we discuss a position restraining scheme during capsid equilibration. With *ω* = 0.7, we ensure that position restraints are only applied to peripheral proteins.

#### 2.1.3 *Step 3:* Position Restraint Generation

The global topology of the capsid surface must not distort while the solvent equilibrates. To this end, we place the capsid surface atop an implicit wall and apply scaled position restraints (Fig. 1c). The wall is constructed using the GROMACS *Walls* function and is placed at *z* = 0 and *z* = *h_box_*, where *h_box_* is the height of the simulation box in the *z* dimension. We use the CHARMM36 *C_δ_* parameters as the wall atomtype, with a number density of 5 atoms/nm^2^, to prevent diffusion through the wall. Periodic boundaries are applied in the *x* and *y* dimensions such that the wall is infinitely repeating perpendicular to the capsid surface normal. We use a stiff 10-4 Lennard-Jones potential integrated over the wall surface. During energy minimization, we apply a linear potential forcing atoms to a separation of 0.1 nm from the wall. This forces the positions of all atoms in the system between *r_z_ >* 0 and *r_z_ < h_box_*.

Capsid atoms near the wall (*r_z_ <* 1.5 nm) have their positions restrained according to a harmonic potential (*K* = 1000 kJ mol^−1^ nm^−2^), while capsid atoms far from the wall (*r_z_ >* 3.0 nm) are fully flexible (*K* = 0 kJ mol^−1^ nm^−2^). At intermediate distances, force constants are weighted from 0 to 1000 kJ mol^−1^ nm^−2^ according to an inverse sigmoid function:

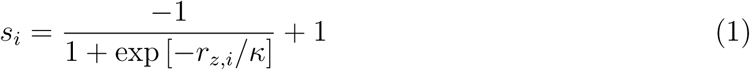

where *r_z,i_* is the *z*-coordinate of capsid atom *i* and *κ* controls the rate of change of the inverse sigmoid. Force constants are then applied to harmonic restraints of atoms in the intermediate region as *K_i_* = *s_i_K* (Fig. S2a). This allows the ROI to remain fully flexible whereas peripheral capsid regions provide the correct molecular context.

In addition to providing a stable platform for the capsid surface to adsorb to, the implicit wall serves another crucial function. While viral capsids are typically permeable to water and salt ions,^85,86^ larger molecules such as excipients are restricted to the capsid exterior. The wall at *z* = *h_box_* prevents excipients from diffusing across periodic boundaries into the capsid interior. Additionally, we apply flat-bottom position restraints that restricts excipients from volumes within 1.5 nm from the wall at *z* = 0 nm (Fig. S2a), which corresponds to a distance of *r* = 9.5 nm from the initial positions. This restraint prevents excipients from interacting with the fully-restrained portion of the capsid surface (Fig. 1c), and takes the form:

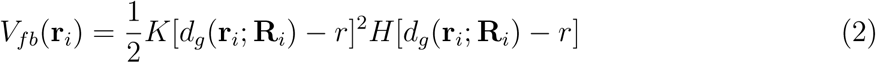

where **R***_i_*is a reference position for excipient atom *i*, **r***_i_*is the position of the heavy atom *i*, *r* is the distance from the center of the flat-bottom potential, *d_g_*(**r***_i_*; **R***_i_*) is the Euclidean distance from the reference position, and *H* is the Heaviside step function.

#### 2.1.4 *Steps 4 and 5:* Restrained and Unrestrained Molecular Dynamics Simulations

Excipient positions are initialized as a 0.1 nm thick slab near the top (*z* = 11 nm) of the simulation box (Fig. S2a). This slab is used as the reference position for the flat-bottom restraint described in Section 2.1.3. Initialization of excipients in this manner prevents excipient placement in the capsid interior. The relative accumulation/depletion of an excipient near the virus surface is described by the preferential interaction coefficient, Γ*_XV_*. This property is calculated in simulations using the two-domain formula^87–89^ given by:

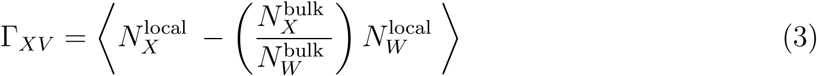

where *V* denotes the viral surface, *X* represents an excipient species, and *W* indicates water. *N* represents the number of molecules of a given species in the local or bulk domain of the viral surface. Equilibrium simulations of the capsid surface model with restraints applied are run for 50 ns to monitor Γ*_XV_* convergence (Fig. 1d). We observe Γ*_XV_* convergence within this simulation time (Fig. S3), consistent with previous studies.^89,90^

All simulations were performed using GROMACS,^91,92^ with versions 2020.4 (for production runs on SDSC Expanse^93^) or 2023.2 (for equilibration runs and nanofragmentation simulations on MSI). Systems were initially subject to energy minimization using the steepest descent algorithm.^94^ An initial NVT equilibration was carried out for 1 ns at 300 K using the V-rescale thermostat,^95^ followed by a 50 ns NVT equilibration using the Nosé-Hoover thermostat.^96^ During equilibration, capsid positions are restrained as described in Section 2.1.3. Following equilibration, production runs were completed for 50-400 ns with capsid harmonic restraints removed (Fig. 1e) but excipient flat-bottom restraints retained. For the 2- and 3-fold surfaces, 100 ns production runs were carried out; for the 5-fold surface without excipients, 400 ns; and for 5-fold surfaces with excipients, 100 ns. We consider our capsid surface simulations to be at equilibrium if (i) ROI capsid protein monomers have steady-state RMSD *C_α_* values varying within a consistent range (Fig. S4), and (ii) excipient preferential interaction coefficients vary around a steady-state average value (Fig. S5).

The Particle Mesh Ewald (PME) algorithm^97^ was used for electrostatic interactions with a cut-off of 1.2 nm. A reciprocal grid of 42 x 42 x 42 cells was used with 4^th^ order B-spline interpolation. We applied a correction to the force and potential in the *z*-dimension to generate a pseudo-2D summation.^98^ A force-switching scheme from 1–1.2 nm was used to describe long-range van der Waals interactions. The neighbor search was performed every 10 steps. Lorentz-Berthelot mixing rules^99,100^ were used to calculate non-bonded interactions between different atom types.

The CHARMM36 force field^101,102^ was used to parameterize virus atoms, arginine (ARG), glutamate (GLU), glycine (GLY), Na^+^, and Cl^−^, while a modified Kirkwood-Buff based parameter set was used for trehalose (TRE) and sorbitol (SOR). ^103^ All systems were solvated with standard TIP3P water^104^ and an NaCl concentration of 0.15 M. Additional Na^+^ and Cl^−^ atoms were added to neutralize the system, as necessary. The protonation states of titratable virus residues at pH 7 were determined by PROPKA3^105^ applied to a PPV monomer (PDB: 1K3V).^84^ The same workflow was used for preparing the fully assembled PPV capsid, albeit without restraints. In totality, for capsid protein monomers, we obtained 5.4 *µ*s of restrained and 12.6 *µ*s of unrestrained PPV capsid protein dynamics from surface systems. 25.5 *µ*s of capsid protein dynamics from the fully assembled capsid were obtained to serve as our ground truth.

Selecting the height of the simulation box, *h_box_*, is a non-trivial task. Too small an *h_box_* and the wall placed at *z* = *h_box_* will interact with the hydration shell of the capsid. Too large an *h_box_* and significant computational cost is incurred. Our criteria for a sufficiently large *h_box_* is that (i) interactions between periodic capsid images should be minimal, and (ii) bulk properties of water can be recovered far from the walls and capsid surface. We address (i) by selecting an *h_box_* value at least 1.5 nm greater than the height of the capsid surface, ensuring periodic capsid surfaces are at a distance greater than the CHARMM non-bonded cut-off distance of 1.0-1.2 nm. For (ii), we ultimately found *h_box_* = 12 nm is the minimum height sufficient to recover several bulk properties of TIP3P water (Figs. S6, S7, and S8).

#### 2.1.5 *Step 6:* Nanofragmentation Simulations

Ghaemi *et al.*^82^ demonstrated that inducing macroscopic cracks across hepatitis B virus (HBV) is an effective strategy for assessing capsid stability. Inspired by this approach, we investigate PPV capsid stability through induced capsid nanofragmentation using the GROMACS pulling code.^91,92^ We apply linear pulling forces along the vectors defined by the center-of-mass of the surface, *X_COM,surf_*, and the center-of-mass of each capsid protein, *X_COM,cp_*. We use umbrella pulling with a pulling rate of 0.001 nm ps^−1^ and a force constant of 1000 kJ mol^−1^ nm^−2^ (see SI Section 4, Fig. S9). For the simulations reported here, we found that the cracks formed within 2.5 ns (determined by visual inspection and force curves) and therefore, the simulations were run for 2.5 ns or until 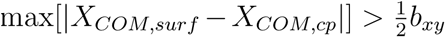, where *b_xy_* is the minimum length of the box in the *x*- and *y*-dimensions. Frames are extracted from pulling simulations every 10 ps for further analysis.

We initialize pulling simulations using configurations from equilibrium MD simulations. Specifically, we extract a frame every 5 ns from the final 25 ns of each production run, resulting in 5 independent configurations for pulling. All data arising from pulling simulations are thus shown as the mean and standard error from 5 such simulations. The stage from which starting configurations are extracted has significant impacts on the outcome of excipient-mediated stability. Configurations taken from unrestrained, equilibrium simulations provide the best agreement with experiment (Fig. S10).

### 2.2 Capsid Dynamics, Structure, and Stability

#### 2.2.1 Dynamic Cross-Correlation

Dynamic cross-correlation (*d_corr_*) analysis is useful in quantifying the correlation coefficients of motions between atoms.^106^ We compute intraprotein correlations between the *C_α_*atoms of residues *i* and *j*, following:

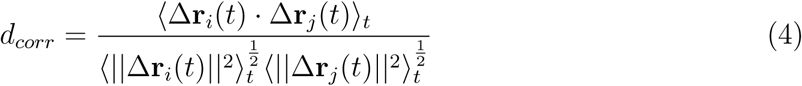

where Δ**r***_i_*(*t*) = **r***_i_*(*t*) − ⟨**r***_i_*(*t*)⟩*_t_* and ⟨·⟩ denotes the time-averaged value. To assess monomer-monomer correlations within virus capsids, we calculate *d_corr_* using the centers-of-mass of capsid proteins.

#### 2.2.2 Residue Interaction Networks

We compute residue interaction networks (RINs) to quantify intraprotein structure. ^107–110^ To construct these graphs, we consider a pair of residues *i, j* to be interacting if the following criteria are met: (i) at least one pair of heavy atoms is within 0.4 nm, and (ii) *i* and *j* are at least 3 residues away in the amino acid sequence. All residues are then assigned to nodes and edges are created between interacting nodes.

To examine the network of intramolecular interactions within capsid protein monomers, we measure betweenness centrality,^111–113^ *C_b_*, of nodes within each RIN. Betweenness centrality quantifies the number of times a node acts as a bridge along the shortest path between two other nodes. This quantity is often thought of as the propensity for a node to act as a “hub” of information transfer. Betweenness centrality is computed as: ^111–113^

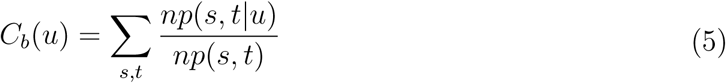

where *np*(*s, t*) is the number of shortest paths between nodes *s* and *t* through the connected network, and *np*(*s, t*|*u*) is the number of such paths that pass through node *u*. We compute this metric for residues in capsid protein monomers, and report *C_b_*(*u*) values averaged across monomers as:

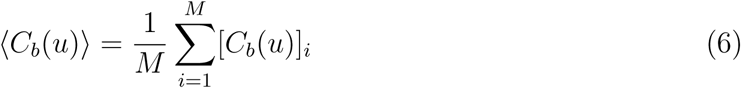

where *u* corresponds to an identical residue in all capsid proteins, *M*.

#### 2.2.3 Sphericity

Capsid sphericity, Ψ, is obtained according to Wadell’s definition:^114^

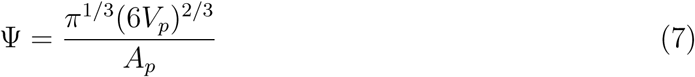

where *V_p_* is the volume of the capsid and *A_p_* is the surface area. Ψ = 1 for spheres, while all non-spheres have Ψ *<* 1. We estimate *V_p_* and *A_p_* from the convex hull defined by the point cloud of capsid atomic coordinates, constructed using Trimesh. ^115^

#### 2.2.4 Residue-Level Capsid Fragmentation

We define the fraction of native interfacial contacts, *Q_IF_*, to monitor fragmentation progression during pulling simulations:

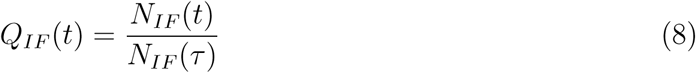

where *N_IF_* is the number of interfacial contacts at time *t* or a reference configuration at *t* = *τ*. We define residues *i* and *j* as a native interfacial contact if the following criteria are met: (i) residues *i* and *j* do not belong to the same capsid protein, and (ii) the minimum distance between heavy atoms, *r_ij_*, is less than 0.45 nm in the reference configuration.

To measure the difference in *Q_IF_* (*t*) curves in excipient and reference solutions, we compute *Q_IF_* (*t*)*^Exc^* − *Q_IF_* (*t*)*^Ref^*. In general, we use the virus in solution with 0.15 M NaCl as our reference solution. We then compute the sum of differences over the pulling trajectory:

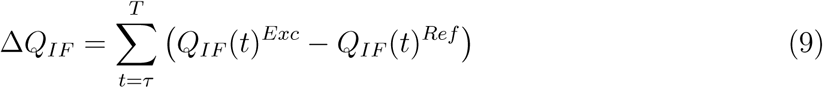

where *T* is the total simulation time. Mean *Q_IF_* values of an individual residue *i* are computed by considering all observed interfacial contacts *ij* involving *i*. Errors are reported as standard errors from the mean of five replicate simulations.

#### 2.2.5 Temperature Stability Assays

Wet-lab experiments were performed to measure the thermal stability of PPV. Liquid samples of PPV in buffer or excipient solutions were prepared in triplicate and were put in either a heat block at 60 ^◦^C^116^ or in a fridge at 4 ^◦^C for 72 hours. The titer of PPV was determined by 3-(4,5-dimethylthiazol-2-yl)-2,5-diphenyltetrazolium bromide (MTT) colorimetric cell viability assay.^117^ Additional experimental details, including materials and methods, are provided in SI Section 7.

## 3 Results and Discussion

### 3.1 Construction and Performance of PPV Surface Models

PPV is a non-enveloped virus with a single stranded DNA genome.^118^ The PPV capsid is an icosahedral shell comprised of 60 copies of viral proteins (VP) VP1, VP2, and VP3 in a 1:10:1 ratio.^84^ Due to the relatively small size and structural simplicity of PPV, recent studies have employed the virus as a model for investigating virus purification and thermostabilization techniques.^116,117,119,120^ PPV, and other similar parvoviruses, are often used in virus removal studies for biotherapeutic manufacturing.^121–123^

The major capsid protein of PPV, VP2, is known to self-assemble into virus-like particles (VLPs) that are non-pathogenic, but are morphologically similar to native PPV virions. ^124^ On the PPV surface, prominent structural regions of interest (ROIs) include a canyon surrounding a pore at the 5-fold axis of symmetry, protrusions located at the 3-fold axis of symmetry, and a dimple at the 2-fold axis of symmetry (Fig. 2a). ^84^ To capture the PPV ROIs, we utilize the CapSACIN workflow to generate PPV surface models centered on the 2-, 3-, and 5-fold axes of symmetry (Fig. 2).

**Figure 2:**
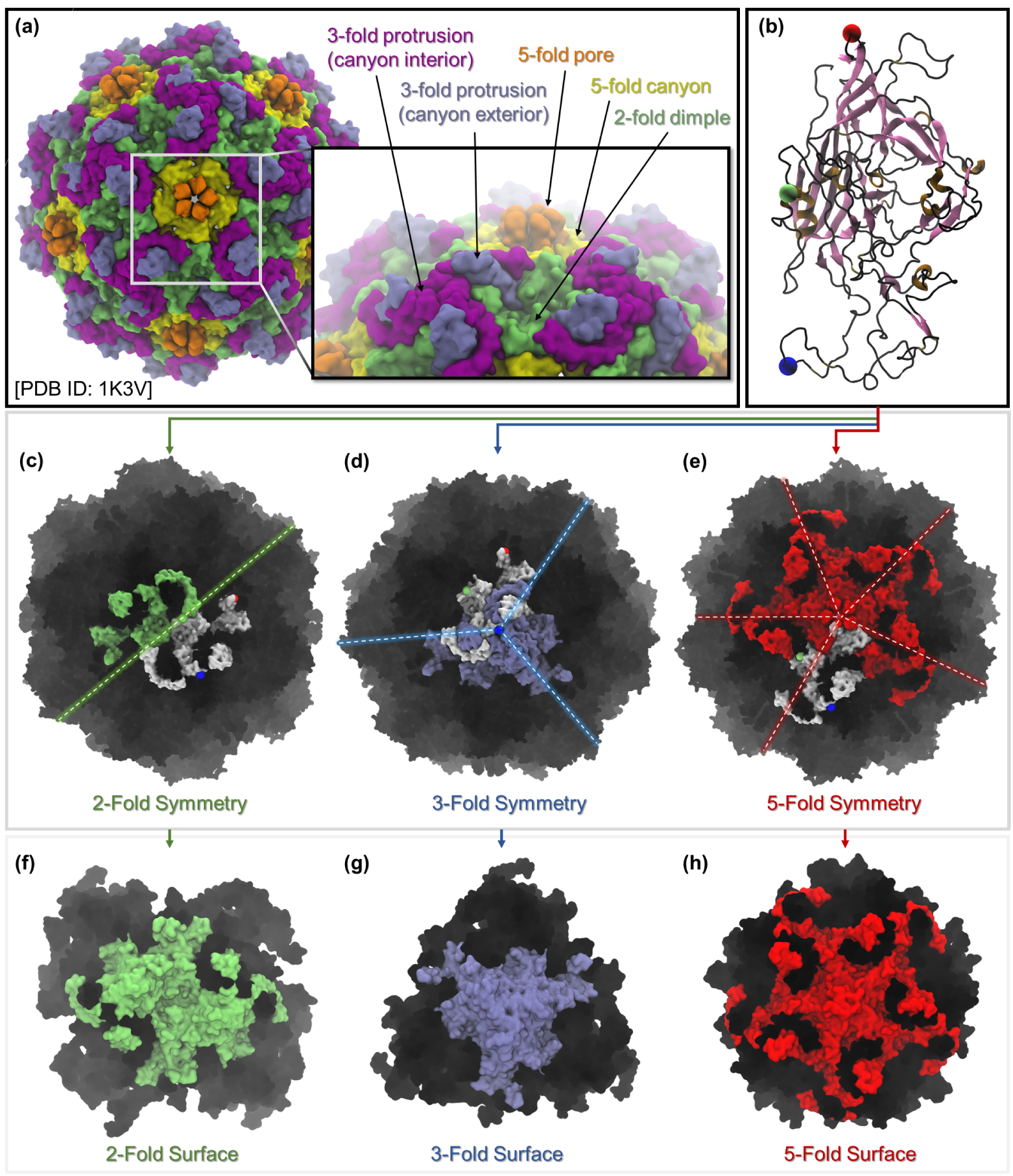
Construction of model PPV surfaces. (a) PPV capsid structure, including key structural ROIs: 5-fold pore (orange), 5-fold canyon (yellow), 3-fold protrusions (blue, purple), and 2-fold dimple (green). (b) PPV monomeric capsid protein, with reference atoms for surfaces shown as spheres (2-fold, green; 3-fold, blue; 5-fold, red). (c) PPV 2-fold axis of symmetry. (d) PPV 3-fold axis of symmetry. (e) PPV 5-fold axis of symmetry. In (c-e), the reference capsid protein from the deposited PDB entry is colored in white, symmetrical proteins are green, blue, or red, and the rest of the capsid is black. Surface models are shown for (f) 2-fold, (g) 3-fold, and (h) 5-fold axes of symmetries. In (f-h), the 2-, 3- and 5-fold ROIs are colored in green, blue, and red, respectively, while peripheral proteins are black.

Each capsid surface model includes a complete ROI (5-fold pore, 3-fold protrusion, or 2-fold dimple), as well as a variable number of peripheral proteins for each surface. The number of peripheral proteins to include depends on the complexity and number of proteins within the ROI, and at the very least, should be sufficiently high such that no ROI atoms are included in the positional restraining scheme. With a slicing weight of *w* = 0.7, each of our PPV surface systems had a 2:1 ratio of peripheral to ROI proteins. The resulting surface systems are between 20–23% the size of a fully assembled capsid system (Table S1). Since the PPV 5-fold surface is the largest ROI (Fig. 2c-h) we studied, we focus the majority of the discussion on this surface. Similar analyses were performed for the other surfaces and are reported in the SI.

The 5-fold surface model consists of nearly 2.5 million fewer atoms relative to the fully-assembled capsid. Atomistic MD simulations for the 5-fold surface are between 5–18x faster than the assembled capsid depending on the available resources and computing hardware (Table 2). We also observe the 5-fold surface scales linearly with CPU cores, while the assembled capsid becomes sublinear as computational resources increase (Fig. S15). The speed-up we observe is comparable to popular coarse-graining strategies,^125,126^ yet the virus surface models presented in this study can be investigated at atomistic resolution.

**Table 2:** Simulation performance and computational cost associated with fully assembled and surface models of PPV. Error estimates of simulation performance, where reported, are the standard deviation from three replicate hsimulations.

| System | System Size<br>[atoms] | CPU Cores | Performance<br>[ns/day] |
| --- | --- | --- | --- |
| Assembled PPV Capsid | 3,146,527 | 128 | <1 |
| | | 256 | $3.881 \pm 0.1$ |
| | | 512 | $7.484 \pm 0.07$ |
| | | 1024 | $12.279 \pm 0.29$ |
|  |  | 2048 | 13.886 |
| PPV 5-Fold Surface | 745,274 | 128 | $11.031 \pm 0.01$ |
| | | 256 | $17.450 \pm 1.56$ |
| | | 512 | $29.537 \pm 0.92$ |
| | | 1024 | $54.188 \pm 0.96$ |
| | | 2048 | $86.031 \pm 11.78$ |
| PPV 3-Fold Surface | 635,743 | 128 | $11.370 \pm 0.56$ |
| | | 256 | $17.417 \pm 2.88$ |
| | | 512 | $34.719 \pm 1.56$ |
| | | 1024 | $54.677 \pm 9.36$ |
| | | 2048 | $72.638 \pm 8.42$ |
| PPV 2-Fold Surface | 692,773 | 128 | $11.033 \pm 0.42$ |
| | | 256 | $18.766 \pm 0.60$ |
| | | 512 | $30.275 \pm 4.73$ |
| | | 1024 | $50.925 \pm 1.36$ |
| | | 2048 | $57.330 \pm 13.34$ |

To highlight the necessity of peripheral, context-providing proteins in simulating capsid ROIs, we compare our surface models to subunit models—which include an ROI, but lack peripheral proteins (Fig. 3d-f). Fig. 3a-c shows the change in the root-mean-square-deviation (RMSD) of *C_α_* atoms taken from ROIs of the assembled capsid, capsid surface models, and capsid subunits. In all cases, the ROI extracted from the first frame of the simulation is used as a reference configuration. These data show that for the assembled capsid (Fig. 3a) and capsid surface models (Fig. 3b), the conformational stability of ROI proteins is high. In subunit models, which lack peripheral proteins, RMSD*_Cα_* varies more over the course of the simulation (Fig. 3c). Such deviations will impact both the conformational ensemble and dynamics of the ROI. In the remainder of this study, we demonstrate that PPV surface models accurately reproduce local and global structural and dynamic features of the ROI, compared to fully assembled capsid ROIs.

**Figure 3:**
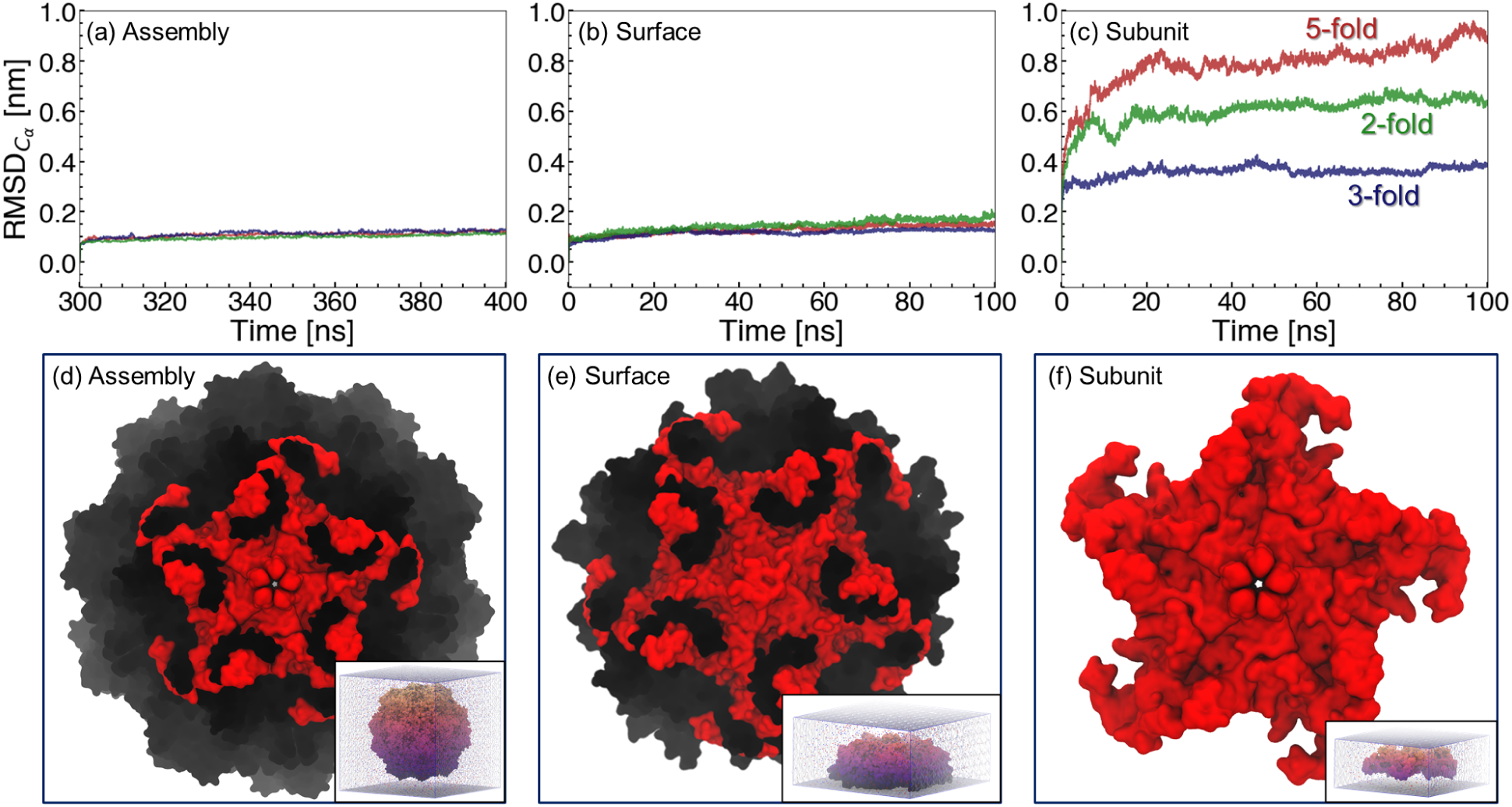
Conformational deviations of capsid ROIs relative to initial configurations, as measured by RMSD*_Cα_*. 100 ns portions of (a) assembled capsid (300-400 ns), (b) surface models, and (c) capsid subunit trajectories are shown. The 5-, 3-, and 2-fold ROIs are colored red, blue, and green, respectively. Representative snapshots are shown for (d) assembled capsid, (e) 5-fold surface model, and (f) 5-fold subunit. Capsids are shown in surface representations from the top-down and are colored according to ROI (red) and peripheral, context-providing proteins (black). The insets in (d-f) show the relative differences in system sizes. Water is shown as a transparent box, while Na^+^ and Cl*^−^* ions are shown as blue and red spheres, respectively.

### 3.2 Intraprotein Structure and Dynamics: Local Similarity

For PPV surface models to accurately reflect the structure and dynamics of a fully assembled capsid, capsid proteins must behave identically at the monomer level between the two representations. In the fully assembled PPV capsid, individual capsid protein monomers have *RMSD_C__α_* values between 0.125–0.24 nm for the majority of the production run (Fig. S4a). In the PPV 5-fold surface model, ROI monomers have *RMSD_C__α_* values within a similar range (0.1-0.215 nm), while peripheral monomers are subject to larger conformational deviations from the initial state (Fig. S4b). To further assess intraprotein dynamics of PPV capsid proteins, we compute dynamic cross-correlations between all residue-residue pairs at the 5-fold axis of symmetry (Fig. 4), as well as for 3- and 2-fold surface models (Fig. S16).

**Figure 4:**
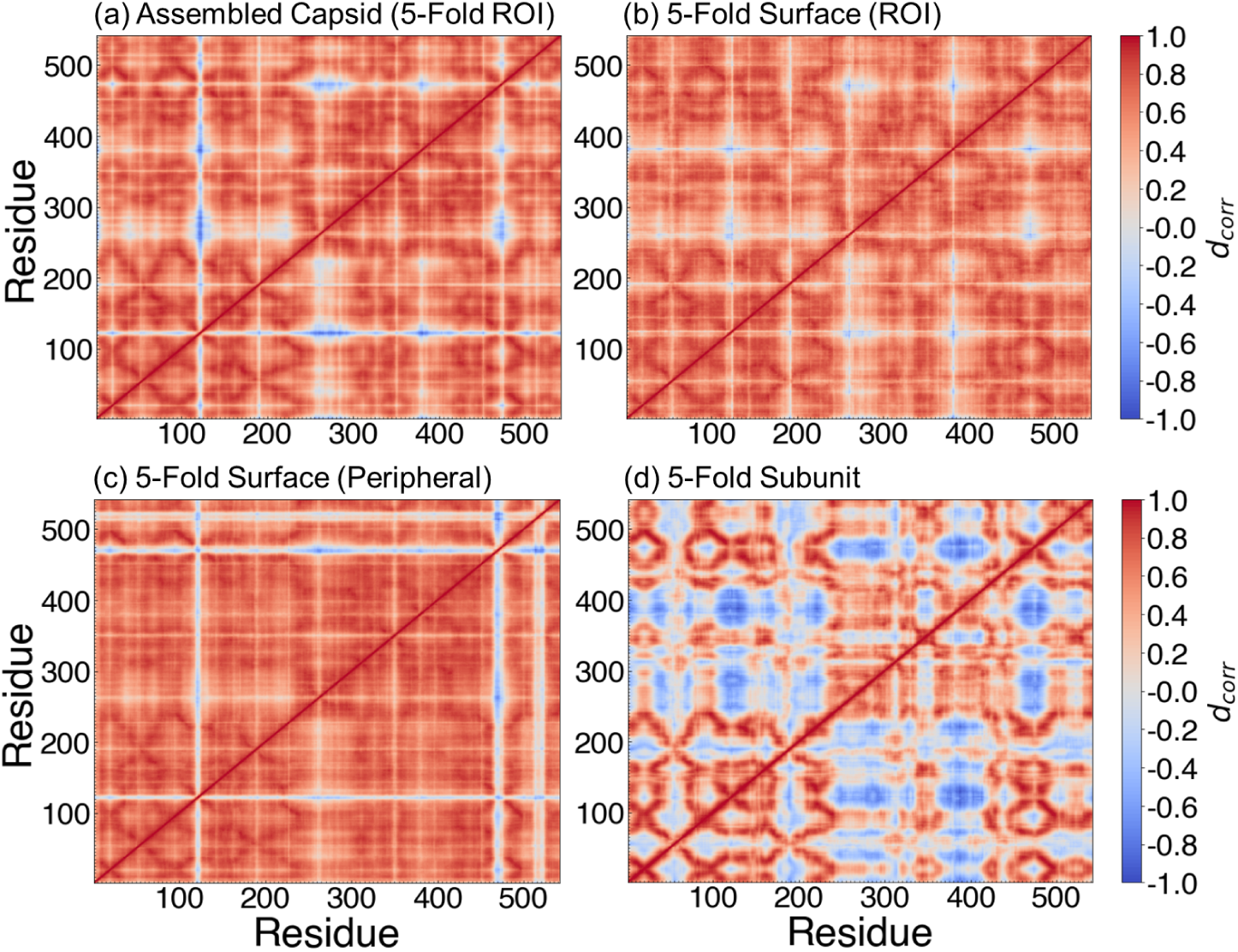
Dynamic cross-correlation matrices of residue-residue pairs within PPV 5-fold capsid proteins. (a) 5-fold ROI subset from the assembled capsid, (b) 5-fold surface ROI, (c) 5-fold surface peripheral proteins, and (d) 5-fold subunit. *d_corr_* values are colored from blue to white to red, with blue indicating strong negative correlation, red indicating strong positive correlation, and white indicating weak or no correlation.

Intraprotein correlations from 5-fold ROI proteins in the assembled capsid (Fig. 4a) and 5-fold surface (Fig. 4b) are in excellent agreement. In these proteins, residues 115–125 are negatively correlated with residues 250–300 and 470–475. These residues roughly correspond to decorrelated motions between the 2-fold dimple (115–125) and stretches of buried residues near the 3-fold protruding spikes (250–300) or near the 5-fold canyon (470–475). Similar results are obtained for the 3- and 2-fold surfaces (Fig. S16), demonstrating that internal dynamics of capsid proteins are preserved even in distinct ROIs of capsid surface models. Greater disagreement with assembled capsid proteins is observed for peripheral proteins of the 5-fold surface (Fig. 4c), where a marked decrease in correlated residue-residue fluctuations is observed from residues ∼510–530 (Fig. 4c). Within the 5-fold subunit, there is a significant increase in negatively correlated motions (Fig. 4d), relative to the assembled capsid and capsid surface models. To quantitatively measure the similarity between these matrices, we compute the inner product angle between matrix pairs according to 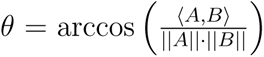, where ⟨*A, B*⟩ is the inner product and ||*A*|| and ||*B*|| are the norms. *cos*(*θ*), then, ranges from −1 to 1, with 1 indicating positive correlation, -1 indicating negative correlation, and values close to 0 indicating no correlation. Compared to the assembled capsid ROI, cos(*θ*) for the 5-fold surface ROI is 0.964, 0.947 for the peripheral 5-fold surface, and 0.689 for the 5-fold subunit. Together, these data demonstrate that including peripheral, context-providing proteins is essential to capture the full intraprotein dynamics within a capsid ROI.

In addition to characterizing intraprotein dynamics, we assess intraprotein structure by constructing residue interaction networks (RINs) for the PPV systems under study (Fig. 5). RINs provide graph-based representations of pairwise residue-residue interaction networks within proteins and have been useful in providing insights into factors that drive protein structure, function, and stability.^107–110^ Within each RIN, we compute betweenness centrality, *C_b_*, as a measure for how often a given residue acts as a bridge connecting all other residues. Several key residues with high *C_b_* values are identified (Fig. 5, Fig. S18): Tyr185, Pro358, Asn370, Arg404, Thr456, and Gln493. These residues consistently have relatively large *C_b_*, regardless of whether they are extracted from the assembled capsid (Fig. 5a) or surface ROI (Fig. 5b). These data indicate that residue interaction pathways are not significantly disrupted in the ROI of surface models.

**Figure 5:**
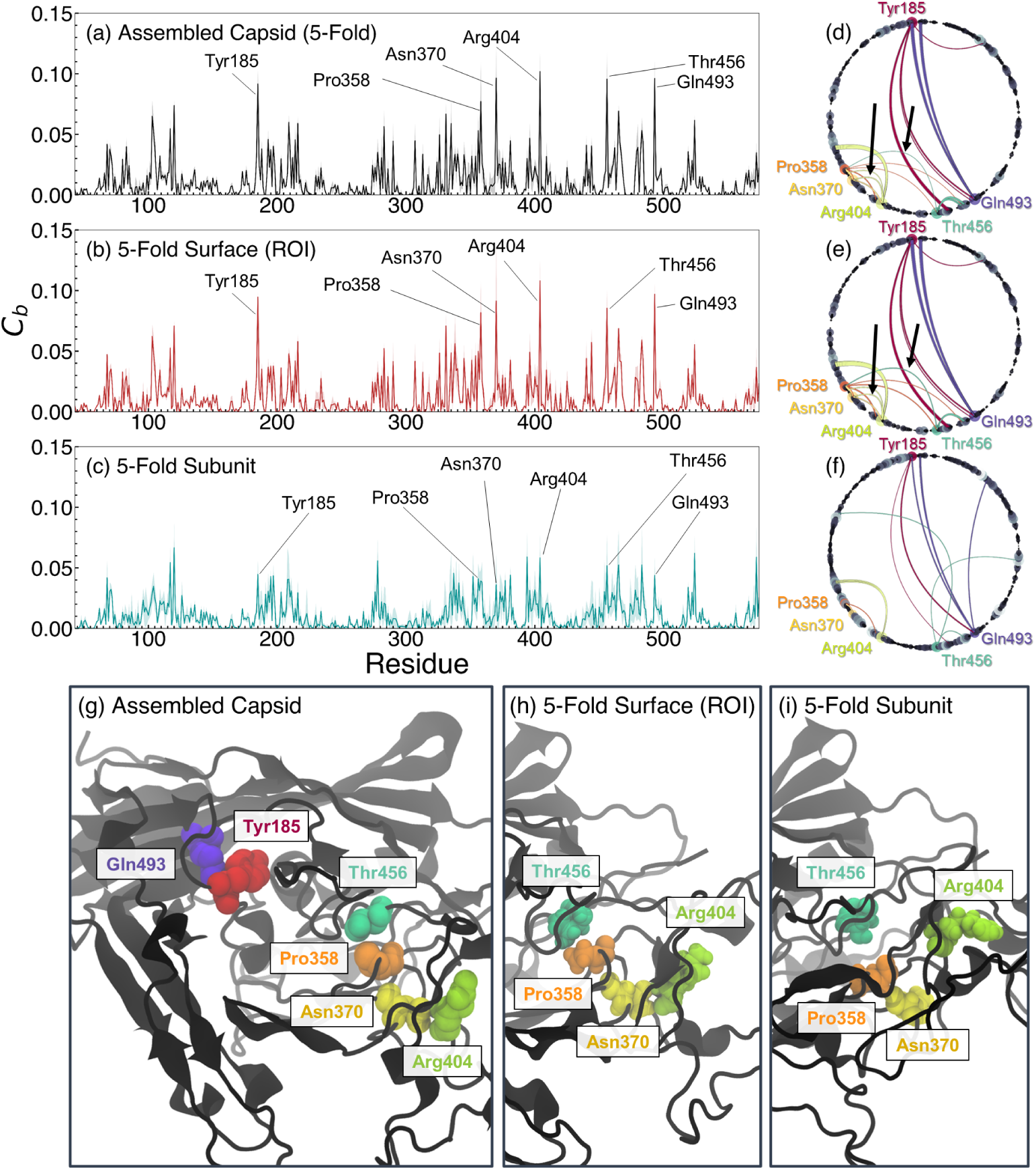
Residue interaction network analysis of PPV capsid proteins. (a-c) Betweenness centrality (*C_b_*) values for capsid protein residues. Arrows point to residues with the highest average *C_b_*value. Standard errors are shown as transparent bars. (d-f) Residue-residue interactions with contact frequency *>*80% during production runs. Edge widths are proportional to the total amount of time two residues are in contact (*r_ij_ <* 0.4 nm). Node sizes are proportional to residue betweenness centrality. Highest betweenness residues are colored as Tyr185, red; Pro358, orange; Asn370, gold; Arg404, yellow-green; Thr456, turquoise; Gln493, violet. Capsid proteins extracted from (a,d) assembled capsid, (b,e) 5-fold surface ROI, and (c,f) 5-fold subunit. (g) Representative configuration showing the relative locations of Tyr185, Pro358, Asn370, Arg404, Thr456, and Gln493. (h) Contacts between Arg404, Asn370, Pro358, and Thr456 in the assembled capsid and 5-fold surface ROI proteins. (i) Broken Asn370–Arg404 contact and Pro358–Thr456 in 5-fold subunit proteins.

In the 5-fold subunit RIN, however, the betweenness centrality of these same residues are reduced (Fig. 5c), implying a change in the connectivity of capsid proteins in the subunit model. Most notably, high-betweenness residues Pro358, Asn370, Arg404, and Thr456 form a persistent network of interactions in the assembled capsid and 5-fold surface ROI (Fig. 5d,e). In the 5-fold subunit, this region is disrupted (Fig. 5f). Specifically, the contacts made between Thr456–Pro358 and Asn370–Arg404 are broken, on average, in 5-fold subunit configurations (Fig. 5g-i). Together, these shifts in centrality highlight how removing the proper molecular context of capsid proteins can give rise to subtle conformational rearrangements, rewire residue interaction networks, and reshape local interaction pathways.

### 3.3 Interprotein Structure and Dynamics: Global Similarity

Dynamic cross-correlations reveal the collective motions of the capsid proteins thereby providing insights into the dynamics of the assembled virus. To quantify these motions, we compute the dynamic cross-correlation of monomer-monomer capsid protein pairs (Fig. 6). The 5-fold ROI (denoted by black boxes in Fig. 6a-c) extracted from the assembled capsid (Fig. 6a) shows strong similarity with the 5-fold surface ROI (Fig. 6b). Within the ROI, proteins exhibit positive correlations. This is consistent with a previous study demonstrating concerted 5-fold motions—such as twists, expansions, and contractions—of another small, icosahedral virus capsid.^127^

**Figure 6:**
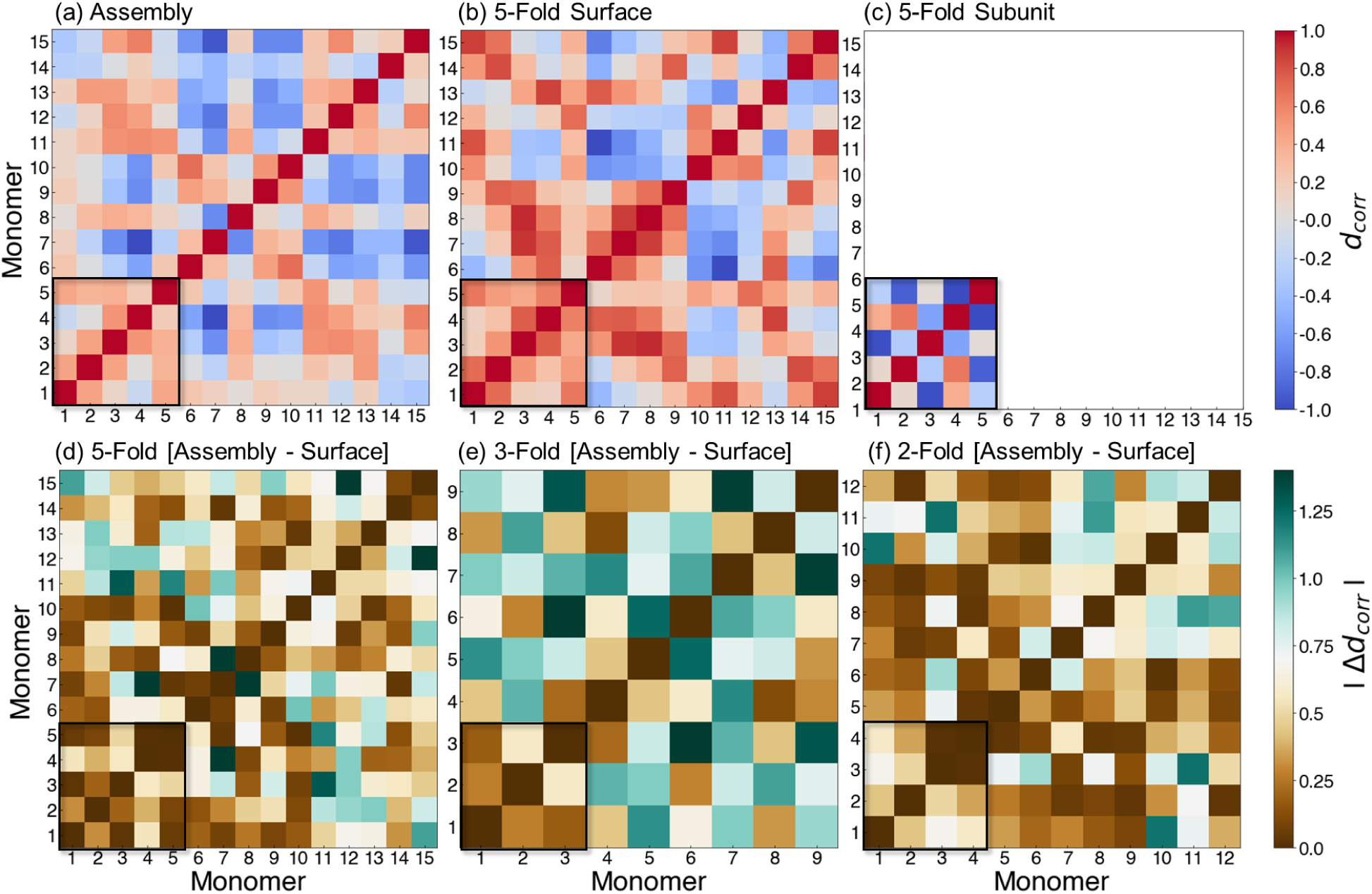
Dynamic cross-correlation matrices of monomer-monomer pairs within PPV capsids. 5-fold capsid proteins taken from (a) the assembled capsid, (b) the 5-fold surface model production run, and (c) the 5-fold subunit. *d_corr_* values are colored from blue to white to red, with blue indicating strong negative correlation, red indicating strong positive correlation, and white indicating weak or no correlation. In (d-f), the magnitude in differences between the assembled capsid and capsid surfaces are plotted. Brown indicates a high degree of similarity, while cyan indicates greater dissimilarity. In all panels, a box is drawn around monomers that form the respective ROI.

The 5-fold subunit (Fig. 6c), on the other hand, fails to accurately capture monomer-monomer correlations present in the assembled capsid. Capsid proteins in the 5-fold subunit are negatively correlated, in stark contrast to the assembled capsid ROI. Peripheral proteins (outside of the black boxes in Fig. 6) are also in poor agreement between the assembled capsid

(Fig. 6a) and the 5-fold surface model (Fig. 6b). In the surface model, 5-fold peripheral proteins show larger positive correlations than the same proteins in the assembled capsid. Correspondingly, cos(*θ*) is 0.955 comparing the assembled capsid 5-fold ROI and surface ROI matrices, 0.324 for the surface peripheral proteins, and 0.280 for the 5-fold subunit. Thus, while peripheral proteins do not capture assembled capsid dynamics, the 5-fold ROI in the surface model more accurately reproduces the 5-fold dynamics of the fully assembled capsid.

Comparable trends emerge for the 3-fold and 2-fold surfaces. In Fig. 6d-f, we plot the difference in correlation strength (|Δ*d_corr_*|) between assembled capsid and capsid surface models. Across all three surfaces, *d_corr_* is similar for ROI proteins in both the capsid surface and assembled capsid, with a few exceptions. At the 5-fold surface, monomer 4 has greater positive correlations with monomers 2 and 3 relative to the assembled capsid (Fig. 6a,b,d); at the 3-fold surface, the correlation between monomers 2 and 3 is more negative (Fig. 6e, Fig. S17a,b); and at the 2-fold surface, monomer 1 has greater negative correlation with monomers 3 and 4 relative to the assembled capsid (Fig. 6f, Fig. S17c,d). These discrepancies may point to altered collective motions at the surface ROIs relative to ROIs from the assembled capsid. However, these differences are minor relative to peripheral, context-providing proteins, where the magnitude of |Δ*d_corr_*| is much larger (Fig. 6d-f). Previous studies have found that out-of-context capsid proteins (e.g., monomers or free pentamers) are more mobile than their assembled counterparts. ^128,129^ Specifically, Pathak and Bandyopadhyay found that residue fluctuations are significantly larger within capsid protein monomers or pentamers, relative to fully assembled minute virus of mice.^128^ Szoverfi and Fejer observed more stable monomers towards the center of cowpea chlorotic mottle virus capsid protein pentamers, while outer monomers are more flexible and exhibit fan-like motions. ^129^ Our observations are consistent with these studies, underscoring the importance of molecular context for using truncated models in virus simulations.

Having established that the 5-fold surface model preserves concerted dynamics of the assembled capsid ROI, we investigated whether the capsid surface model also maintains its overall architecture. In particular, we sought to ensure that no large-scale conformational changes—such as collapse of the surface, distortions in curvature, or unreasonable monomer-monomer separations—arise after removal of position restraints. To this end, we compared the global structure of the 5-fold surface model to the fully assembled PPV capsid using sphericity measurements, Ψ, and monomer-monomer center-of-mass distances, *d_COM_*.

Sphericity measures the deviation of an object from a perfect sphere. For assembled icosahedral viruses, Ψ approaches 1, which we observe for fully assembled PPV (Fig. S19) and Hadden *et al.* observed for HBV. ^64^ For the 5-fold PPV surface model (Fig. 7b), we obtain Ψ values distributed between 0.815–0.825, which is in good agreement with the Ψ values of the 5-fold ROI extracted from the fully assembled PPV capsid (Fig. 7a). *P* (Ψ) is significantly broader for simulations of 5-, 3-, and 2-fold subunits (Fig. 7c), relative to either the assembled capsid or surface models. The 5-fold ROI *P* (Ψ) distributions are centered at 0.822 ± 0.0018 for the assembled capsid, 0.821 ± 0.0019 in the surface model, and 0.840 ± 0.0084 for the capsid subunit. The corresponding values for the 3-fold ROI are 0.892 ± 0.0019, 0.884 ± 0.0020, and 0.893 ± 0.0034; and for the 2-fold ROI the values are 0.879 ± 0.0025, 0.880 ± 0.0018, and 0.886 ± 0.0059. Thus, *P* (Ψ) distributions are consistent between surface models and fully assembled PPV (Fig. 7a,b), with an exception of the 3-fold surface model, which has *P* (Ψ) shifted towards a less spherical shape. *P* (Ψ) is shifted towards broader and more spherical distributions for capsid subunits, reflecting poor preservation of the ROI shape in the absence of peripheral capsid proteins.

**Figure 7:**
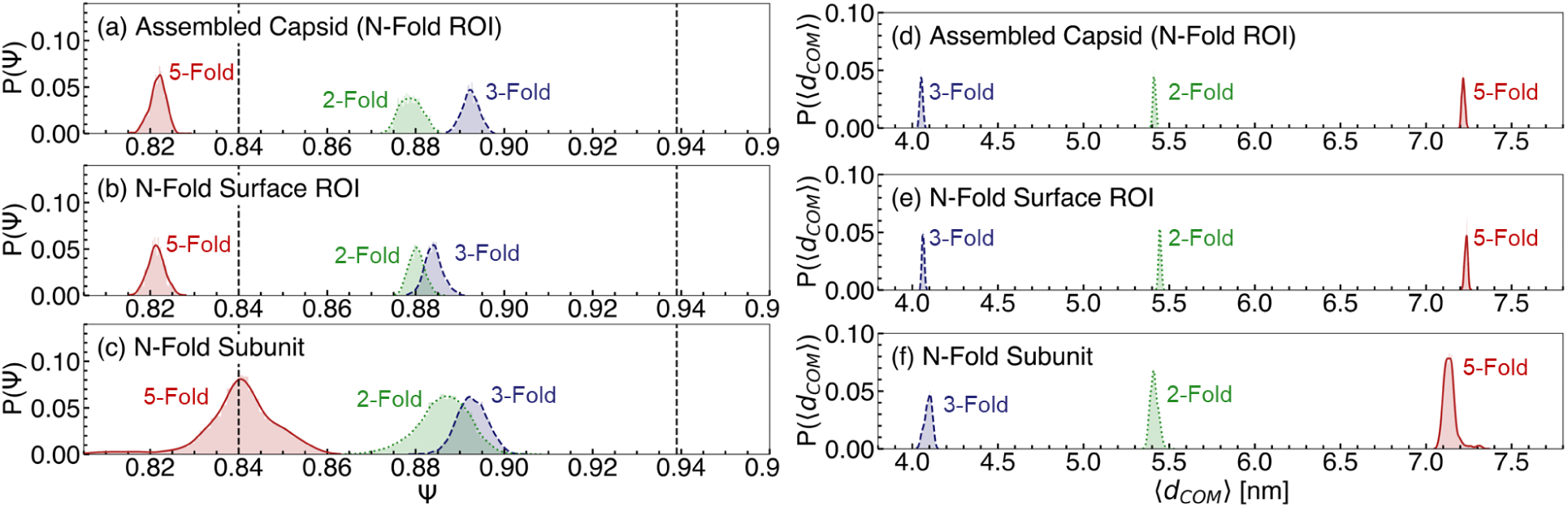
Global structure of the 5-, 3-, and 2-fold ROIs from the assembled capsid, capsid surface models, and capsid subunits. Probability distributions of sphericity, Ψ, are shown for (a) assembled capsid ROIs, (b) capsid surface ROIs, and (c) capsid subunits. Probability distribution of monomer-monomer center-of-mass distances, ⟨*d_COM_* ⟩, are also shown for (d) assembled capsid ROIs, (e) capsid surface ROIs, and (f) capsid subunits. The 5-fold ROI is denoted by a solid red line, the 3-fold ROI by a dashed blue line, and the 2-fold ROI by a dotted green line.

We measure monomer-monomer arrangement using the center-of-mass distance between all protein pairs. For each frame, we compute an average distance ⟨*d_COM_* ⟩, with *P* (⟨*d_COM_* ⟩) shown in Fig. 7d-f. *P* (⟨*d_COM_* ⟩) distributions agree closely between assembled capsid and surface ROIs: The 5-fold, 3-fold, and 2-fold ROI distributions in the fully assembled capsid are centered at 7.2183 ± 0.0088 nm, 4.0531 ± 0.0087 nm and 5.4141 ± 0.0086 nm, respectively. In the surface models, these values are 7.2350 ± 0.0088 nm, 4.0637 ± 0.0071 nm and 5.4466 ± 0.0078 nm. Capsid subunits, meanwhile, have *P* (⟨*d_COM_* ⟩) distributions that reflect a more separated 3-fold ROI and more compact 2- and 5-fold ROIs than the assembled capsid. For the capsid subunits, the 5-fold, 3-fold, and 2-fold ROI distributions are centered at 7.139 ± 0.0401 nm, 4.0933 ± 0.0222 nm and 5.4134 ± 0.0233 nm. These data confirm that truncating the PPV capsid into surface models consisting of both an ROI and peripheral, context-providing proteins is sufficient to retain the overall architecture of the assembled capsid in the ROI. Without peripheral proteins, the ROI can significantly change shape in a manner that does not reflect the properties of the assembled capsid.

### 3.4 Nanofragmentation Along Capsid Protein Interfaces

Non-equilibrium simulations have recently been used to explore the mechanical stability of virus capsids,^82,130–132^ an attractive approach due to direct comparison with experimental methods such as nanoindentation or atomic force microscopy.^133–135^ Inspired by this, we utilize pulling simulations supported by GROMACS to induce nanofragmentation in the three capsid surface models under study. Using this approach, we identify weak interfaces in the PPV capsid and explore how excipient molecules modulate the resistance/susceptibility of PPV surface fragmentation.

We first carried out nanofragmentation simulations of the 5-, 3-, and 2-fold PPV surface models in 0.15 M NaCl solution (Fig. 8a). At the 2-fold surface, the onset of crack formation begins almost immediately, and the surface has a large surface-spanning crack within ∼1.4 ns. The 3- and 5-fold surfaces are more resistant to fragmentation, with the rate of crack formation significantly increasing after ∼1.8 ns and surface-spanning cracks forming by 2.5 ns. By the end of the simulation, the 3-fold surface undergoes slightly more extensive fragmentation than the 5-fold surface. Together, these data point to the 5-fold surface as most resistant to fragmentation, followed by the 3-fold surface, while the 2-fold surface is the most susceptible to fragmentation. In other words, a logical progression of capsid disassembly may proceed as follows: (i) pentamer-pentamer interactions at the 2-fold axis of symmetry break apart, resulting in trimers of pentamers, (ii) pentamer-pentamer contacts begin to separate, leaving behind individual pentamers, which (iii) are the most stable intermediate unit. Such a view supports hierarchical assembly pathways of simple icosahedral viruses wherein pentamers or hexamers form early in the process and nucleate assembly.^136,137^ A susceptible 2-fold interface is consistent with several experimental studies that point to weak 2-fold interactions in T=1 icosahedral viruses, ^138–142^ such as PPV. Meanwhile, a stable 3-fold interface supports a previous study that found certain intertrimer mutations in a closely related parvovirus causes assembly incompetence. ^143^ Molecular Mechanics / Generalized Born

**Figure 8:**
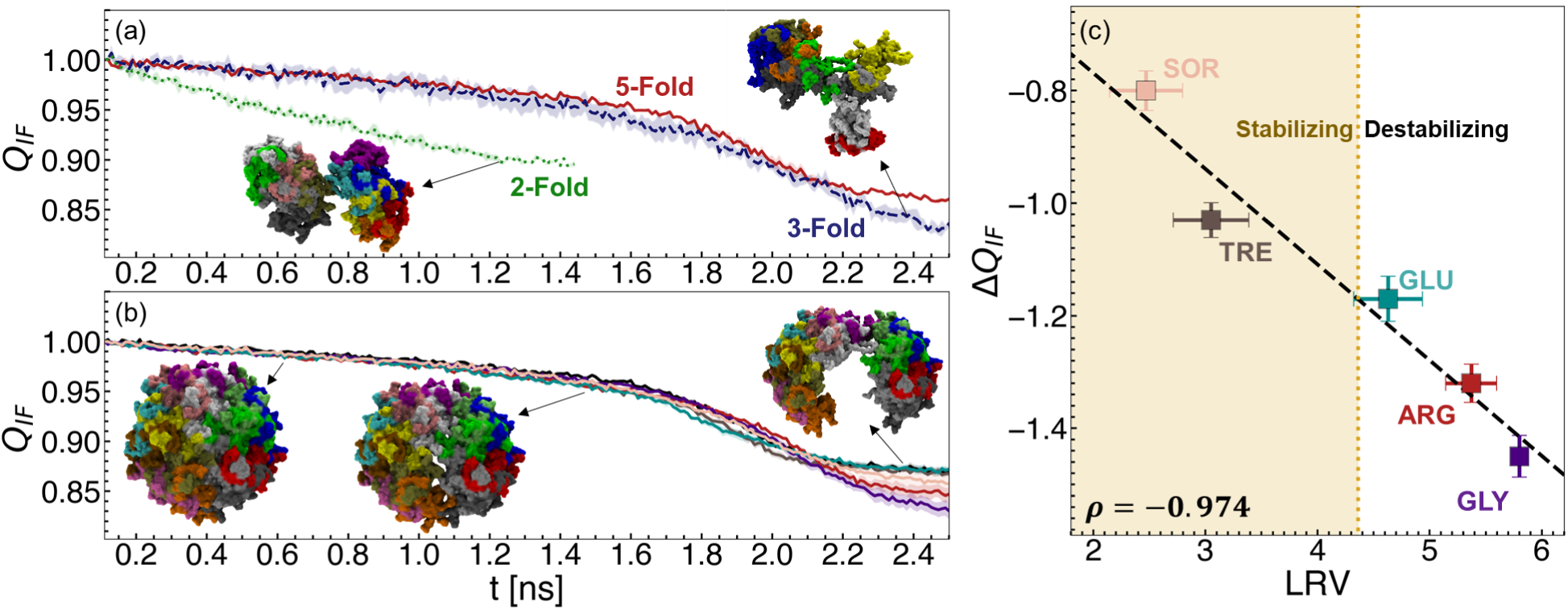
Change in the fraction of native interfacial contacts (*Q_IF_*) during nanofragmentation simulations of PPV 5-fold, 3-fold, and 2-fold surface models. (a) Change in *Q_IF_* for different surface models in 0.15 M NaCl. Colors indicate 5-fold, red; 3-fold, blue; and 2-fold, green. (b) Change in *Q_IF_* for the PPV 5-fold surface in 0.1 M sorbitol (SOR; pink), 0.1 M trehalose (TRE; brown), 0.1 M glutamate (GLU; cyan), 0.1 M arginine (ARG; red), and 0.1 M glycine (GLY; purple). In (a) and (b), representative snapshots of 2-, 3-, and 5-fold surface fragmentation are shown, with arrows indicating the approximate points along the fragmentation curve where snapshots were taken. (c) Correlation between the sum of 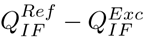, reported as Δ*Q_IF_*, and LRV for excipient solutions. Excipients found to be stabilizing experimentally (LRV*^Exc^ <* LRV*^Ref^*) are highlighted by the yellow shaded region, and the vertical dashed line denotes the LRV obtained following heat treatment of PPV in buffer. The Pearson correlation coefficient, *ρ*, is shown in the bottom left of the panel.

Surface Area (MM-GBSA) calculations were carried out using the HawkDock webserver^144^ to estimate the binding free energy of capsid protein interfaces. These calculations were carried out as a measure of thermodynamic monomer-monomer stability independent from the pulling simulations. Dimers were selected from interacting pairs of capsid proteins from the 5-fold, 3-fold, and 2-fold surface models. The binding free energies were computed for five configurations per fold, with dimers extracted from the same input configurations used for nanofragmentation simulations. MM-GBSA similarly identified the 2-fold interface as the weakest thermodynamically (-167.60 ± 3.425 kcal/mol), followed by the 5-fold interface (-185.10 ± 3.622 kcal/mol), while the 3-fold interface (-616.12 ± 2.324) was most stable through this calculation.

### 3.5 Excipient Effects on PPV Fragmentation are Consistent with Experimental Stability Data

We carried out similar nanofragmentation simulations for 0.1 M solutions of arginine (ARG), glutamate (GLU), glycine (GLY), trehalose (TRE), or sorbitol (SOR) with the 5-fold PPV surface model under study. These excipients were selected due to their prominence in many biological formulations. ^145–148^ Based on the results shown in Fig. 8a, we hypothesized that nanofragmentation of the 5-fold surface should be the rate-limiting step for total capsid disassembly.

Changes in *Q_IF_* during nanofragmentation simulations in excipient solutions are shown in Fig. 8b. Significant deviations in *Q_IF_*do not begin to occur until ∼1.4 ns. Some excipient solutions, including ARG and GLY, delay the rate of fragmentation from ∼1.4–1.8 ns, but ultimately destroy more native interfacial contacts from ∼2.0–2.5 ns than any other excipient. Others, like TRE and GLU, form larger cracks early in the simulation (∼1.4–2.0 ns) but result in roughly the same amount of broken interfacial contacts by the end of the trajectory as in water. SOR results in a *Q_IF_* curve that is nearly overlapping with the behavior of PPV in 0.15 M NaCl for the duration of the simulation.

Experimentally, we determine excipient effects on PPV stability through temperature stability assays. For PPV, we completed an infectivity assay (see SI Section 7) following virus incubation at 60 ^◦^C for 3 days – sufficiently long to observe a significant decrease in the infectious titer.^116^ Log reduction values (LRV) describe the decrease in the infectious titer of heat-treated virus relative to the initial virus solution (Fig. 8c). Lower LRV relative to buffer indicates more infectious virus is present following heat treatment, indicating stability. We found that SOR (2.47 ± 0.326 LRV) and TRE (3.05 ± 0.337 LRV) are stabilizing relative to the reference buffer solution (4.36 ± 0.260 LRV), while GLU (4.63 ± 0.307 LRV), ARG (5.37 ± 0.226 LRV), and GLY (5.8 ± 0.010 LRV) are destabilizing at elevated temperatures.

We use Δ*Q_IF_* as a measure of excipient-mediated stability in nanofragmentation simulations, which describes the cumulative differences in 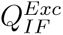 relative to 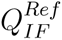. The more negative Δ*Q_IF_*, the more susceptible the surface is to fragmentation in excipient solution relative to in 0.15 M NaCl. All excipient solutions result in Δ*Q_IF_* < 0 (Fig. 8c), implying the PPV 5-fold surface is more susceptible to fragmentation in excipient solutions than in 0.15 M NaCl. While this is at odds with results from heat treatment of PPV experimentally, the rank-ordering of excipients according to Δ*Q_IF_*values is in excellent agreement with experimental LRV data (Fig. 8c). Specifically, SOR (Δ*Q* = −0.80 ± 0.035) and TRE (Δ*Q* = −1.03 ± 0.031) result in the smallest changes to *Q_IF_* relative to in 0.15 M NaCl, GLU (Δ*Q* = −1.17 ± 0.036) and ARG (Δ*Q* = −1.32 ± 0.034) are the next most destabilizing excipients, and GLY (Δ*Q* = −1.45 ± 0.037) is the most destabilizing excipient in simulations. The strong correlation (Pearson correlation coefficient, *ρ* = −0.974 ± 0.0226) between change in stability from experiment and simulation gives us confidence that our approach can effectively identify stabilizing/destabilizing excipients.

## 4 Potential Applications and Limitations of the CapSACIN Framework

The CapSACIN workflow is designed to construct minimal atomistic representations of viral capsids in explicit solvent while retaining important monomer-monomer interactions and dynamics. While we originally developed this method for investigation of capsid-excipient interactions, this framework can be adapted to study interactions between viral capsids and host cell receptors or small molecules. Virus-receptor interaction is a crucial step in infectivity, regulating host range, tissue tropism, and pathogenesis.^149^ Simulations of complex virus-receptor systems are typically performed at the coarse-grained level and can identify large-scale insights such as membrane curvature, binding poses, and global conformational changes.^150,151^ However, molecular-level mechanisms that drive such molecular recognition processes, such as frustration of the local solvent structure, ^152^ hydrogen bonding,^153^ and cation-*π* interactions^154^ are only observable at atomistic resolution. By building models that focus on an ROI hypothesized to interact with a host receptor, the CapSACIN workflow can be leveraged to explore these systems at the level of detail required to investigate the molecular underpinnings of capsid-receptor interactions.

Another potential application of CapSACIN is in the design of capsid assembly modulators (CAMs). CAMs are small molecules that interact with capsid proteins, interfering with formation of the assembled capsid. ^155^ CAM discovery and optimization is accelerated via computer-aided drug design,^156^ leveraging tools such as molecular docking, quantitative structure-activity relationships, and molecular dynamics simulations. Understanding the role of dynamics is especially important in the elucidation of CAM mechanisms, as these molecules have been shown to influence both capsid protein dynamics and conformational states.^157–159^ The current understanding of CAM-mediated changes in capsid dynamics is typically limited to the subunit level due to the computational cost associated with simulating multiple CAM/capsid pairs. CapSACIN provides a means for investigating the CAM-mediated changes in ROI dynamics and conformational states in a computationally-feasible manner, opening the door for high-throughput CAM screening through simulations.

While the CapSACIN workflow can be used to investigate capsid-excipient, capsid-receptor, and capsid-small molecule interactions, our approach is also limited in certain areas. First, our methodology has not yet been adapted for non-icosahedral capsid architectures. Viruses vary in morphology, predominantly taking filamentous, spherical, or pleomorphic shapes. ^160^ Spherical viruses have an icosahedral symmetry, which CapSACIN leverages to construct surface models. Filamentous viruses are rod-shaped and take on a helical symmetry, which would impose a fundamentally different set of geometric rules for building surface models. The second key limitation is that long-range, concerted motions between multiple ROIs are not present in these capsid surface models. Third, representing capsid ROIs as surfaces results in a system that is larger in size than alternative approaches, such as using rotational boundary symmetry conditions (RSBC).^161–163^ However, RSBC is not supported in many mainstream MD engines, whereas CapSACIN is program-agnostic, allowing ROIs to be simulated in any standard MD environment. Finally, CapSACIN in its present form does not accommodate enveloped viruses, whose membrane–protein architectures require an additional modeling layer beyond capsid geometry.

We also note that the nanofragmentation aspect of the CapSACIN workflow is sensitive to both parameter selection (Figs. S9, S11) and the choice of initial configuration (Fig. S10). A systematic analysis on the choice of pulling parameters may be required to generalize to other capsid systems. Given this sensitivity, inferences regarding the stability of capsid surfaces may also differ with different pulling vectors. In the present study, we used pulling vectors between the center-of-mass of the entire surface and the centers-of-mass of individual monomers. However, alternative approaches that, for example, identify collective modes of capsid dynamics or apply a purely radial potential may also serve as a suitable pulling coordinates. In the current approach, we also lack the resolution necessary to decouple mechanical stability and pulling artifacts from purely thermodynamic contributions to monomer-monomer interfacial stability. While our stability results are in good agreement with experimental data, further testing must be done in order to ensure generalizability to new systems.

## 5 Conclusions

In the development of stable viral biologics, excipient selection is a critical step. This stage of development, however, is generally time-consuming and expensive due to the lack of a fundamental understanding of excipient–virus interactions. We have tackled this issue by developing a high-throughput, high-detail workflow—CapSACIN—suitable for examining excipient–virus interactions through atomistic molecular dynamics simulations.

In the present study, we have shown that this method is capable of generating surface models that capture any region of interest from a viral capsid, such as canyons, protrusions, or dimples. We demonstrated that for PPV, inter- and intraprotein structure and dynamics are similar between a fully assembled PPV capsid model and surface models representing the 5-, 3-, and 2-fold axes of symmetries. This approach enables efficient simulations of viral systems when a region of interest is known, resulting in a ∼5× speed-up relative to fully assembled non-enveloped virus capsids and VLPs.

Further, we have shown that by applying radial pulling forces to induce nanofragmentation, we are able to identify weak protein–protein interfaces that are consistent with experimental data. Specifically, we showed that the 2-fold axis of symmetry is the capsid region most prone to nanofragmentation, while the 5-fold and 3-fold regions are more resistant. Through simulations, we are also able to elucidate the effects of excipients on capsid stability. Our simulations reveal a rank-ordering of excipient effects on PPV stability that matches experimental data. Specifically, sorbitol and trehalose are the most stabilizing excipients of PPV, followed by glutamate and arginine, while glycine is the most destabilizing excipient. While the results presented here are consistent with experimental measurements, the rank ordering of excipients exhibits sensitivity to the pulling rate. Systematic evaluation of this dependence will be important to ensure robust determination of excipient-mediated stability across distinct viral systems. In addition, there remains scope for investigating alternative collective variables that may further refine the thermodynamic description.

Qualitatively, our simulations agree with several previous studies that identified trehalose as a broad virus stabilizer,^50,145,148,164–166^ while arginine is capable of viral inactivation^167–169^ and instability.^170^ Similar to PPV, sorbitol has been shown to be a highly effective stabilizer of a live attenuated *Salmonella* Enteriditis vaccine.^171^ On the other hand, sorbitol was not an effective stabilizer of live attenuated influenza vaccine. ^172^ The dependence of virus identity on excipient-mediated effects underscores the need for a reliable approach to predict excipient behavior prior to expending costly experimental resources.

Overall, we present a method for high-throughput investigations into excipient–virus interactions at the atomistic level. The level of detail provided by molecular simulations has the potential to enable viral biologics development from a more information-driven approach, reducing both the time and cost associated with excipient selection. Importantly, by capturing the essential dynamics of virus-like particles without requiring the simulation of entire capsids, our approach provides a practical and scalable strategy for accelerating VLP-based vaccine and gene therapy development. Because VLPs are increasingly used as safe and versatile delivery vehicles, improving their stability through rational excipient selection directly supports broader deployment of next-generation biologics. Our approach further enables future studies aiming to explore different capsid architectures (i.e., icosahedral vs. helical capsid symmetry, T>1 triangulation numbers, or empty vs. full capsids), an expanded library of excipients (i.e., surfactants, peptoids, or oligosaccharides), or exploring different denaturation mechanisms (i.e., temperature, pressure, or chemical denaturation of capsid proteins). Additionally, the generalizability of our approach to viruses that are closely related to porcine parvovirus (i.e., adeno-associated viruses, canine parvovirus, or minute virus of mice) is a clear next step for this work.

## Author Contributions

J.W.P.Z.: Investigation - Computational, Conceptualization, Validation, Formal Analysis, Visualization, Writing - Original Draft, Writing - Review and Editing; I.T.: Investigation - Experimental, Writing - Review and Editing; P.M.: Conceptualization; C.L.H.: Conceptualization, Funding Acquisition, Resources, Writing - Review and Editing; S.L.P.: Conceptualization, Funding Acquisition, Resources, Writing - Review and Editing; S.S.: Conceptualization, Methodology, Validation, Supervision, Project Administration, Funding Acquisition, Resources, Writing - Review and Editing

## Data availability

Data for this article, including simulation parameter files and analysis scripts, and the complete CapSACIN workflow are available at https://github.com/SAMPEL-Group/CapSACIN.

## Acknowledgments

This work is supported by U.S. National Science Foundation DMREF grants #2118788, #2118693, and #2325392. Computational resources were provided by the Minnesota Super-computing Institute at the University of Minnesota - Twin Cities. This work additionally used computing resources from SDSC Expanse at the University of California - San Diego through allocation CHE240118 from the Advanced Cyberinfrastructure Coordination Ecosystem: Services & Support (ACCESS) program, which is supported by U.S. National Science Foundation grants #2138259, #2138286, #2138307, #2137603, and #2138296. J. W. P. Z. acknowledges funding support from the University of Minnesota Graduate School Doctoral Dissertation Fellowship.

## Supporting information

Supporting information: details on surface construction of additional capsids, determination of an appropriate box height, additional simulation parameters, convergence checks, and additional computational details.

## References

(1) D’Amico, C.; Fontana, F.; Cheng, R.; Santos, H. A. Development of vaccine formulations: past, present, and future. Drug Deliv. and Transl. Res. 2021, 11, 353–372.

(2) Zhang, C.; Maruggi, G.; Shan, H.; Li, J. Advances in mRNA Vaccines for Infectious Diseases. Front. Immunol. 2019, 10, 594.

(3) Minor, P. D. Live attenuated vaccines: Historical successes and current challenges. Virology 2015, 479–480, 379–392.

(4) Rappuoli, R.; De Gregorio, E.; Costantino, P. On the mechanisms of conjugate vaccines. Proc. Natl. Acad. Sci. U.S.A. 2019, 116, 14–16.

(5) Bull, J. J.; Nuismer, S. L.; Antia, R. Recombinant vector vaccine evolution. PLoS Comput Biol 2019, 15, e1006857.

(6) Mettu, R.; Chen, C.-Y.; Wu, C.-Y. Synthetic carbohydrate-based vaccines: challenges and opportunities. J Biomed Sci 2020, 27, 9.

(7) Delany, I.; Rappuoli, R.; De Gregorio, E. Vaccines for the 21st century. EMBO Mol Med 2014, 6, 708–720.

(8) Aljabali, A. A.; Hassan, S. S.; Pabari, R. M.; Shahcheraghi, S. H.; Mishra, V.; Charbe, N. B.; Chellappan, D. K.; Dureja, H.; Gupta, G.; Almutary, A. G. et al. The Viral Capsid As Novel Nanomaterials for Drug Delivery. Future Sci. OA 2021, 7, FSO744.

(9) Comisel, R.-M.; Kara, B.; Fiesser, F. H.; Farid, S. S. Lentiviral vector bioprocess economics for cell and gene therapy commercialization. Biochemical Engineering Journal 2021, 167, 107868.

(10) Xiang, Y. S.; Hao, G. G. Biophysical characterization of adeno-associated virus capsid through the viral transduction life cycle. Journal of Genetic Engineering and Biotechnology 2023, 21, 62.

(11) Eid, F.-E.; Chen, A. T.; Chan, K. Y.; Huang, Q.; Zheng, Q.; Tobey, I. G.; Pacouret, S.; Brauer, P. P.; Keyes, C.; Powell, M. et al. Systematic multi-trait AAV capsid engineering for efficient gene delivery. Nat Commun 2024, 15, 6602.

(12) Carneiro, A. D.; Schaffer, D. V. Engineering novel adeno-associated viruses (AAVs) for improved delivery in the nervous system. Current Opinion in Chemical Biology 2024, 83, 102532.

(13) Nisanov, A. M.; Rivera De Jesús, J. A.; Schaffer, D. V. Advances in AAV capsid engineering: Integrating rational design, directed evolution and machine learning. Molecular Therapy 2025, 33, 1937–1945.

(14) Lua, L. H.; Connors, N. K.; Sainsbury, F.; Chuan, Y. P.; Wibowo, N.; Middelberg, A. P. Bioengineering virus-like particles as vaccines. Biotech & Bioengineering 2014, 111, 425–440.

(15) McNeale, D.; Dashti, N.; Cheah, L. C.; Sainsbury, F. Protein cargo encapsulation by virus-like particles: Strategies and applications. WIREs Nanomed Nanobiotechnol 2023, 15, e1869.

(16) Dashti, N. H.; Abidin, R. S.; Sainsbury, F. Programmable *In Vitro* Coencapsidation of Guest Proteins for Intracellular Delivery by Virus-like Particles. ACS Nano 2018, 12, 4615–4623.

(17) Das, S.; Zhao, L.; Crooke, S. N.; Tran, L.; Bhattacharya, S.; Gaucher, E. A.; Finn, M. G. Stabilization of Near-Infrared Fluorescent Proteins by Packaging in Virus-like Particles. Biomacromolecules 2020, 21, 2432–2439.

(18) Wilkerson, J. W.; Yang, S.-O.; Funk, P. J.; Stanley, S. K.; Bundy, B. C. Nanoreactors: Strategies to encapsulate enzyme biocatalysts in virus-like particles. New Biotechnology 2018, 44, 59–63.

(19) Jordan, P. C.; Patterson, D. P.; Saboda, K. N.; Edwards, E. J.; Miettinen, H. M.; Basu, G.; Thielges, M. C.; Douglas, T. Self-assembling biomolecular catalysts for hydrogen production. Nature Chem 2016, 8, 179–185.

(20) Wang, Y.; Uchida, M.; Waghwani, H. K.; Douglas, T. Synthetic Virus-like Particles for Glutathione Biosynthesis. ACS Synth. Biol. 2020, 9, 3298–3310.

(21) Organization, W. H. Temperature sensitivity of vaccines. 2006.

(22) Hanson, C. M.; George, A. M.; Sawadogo, A.; Schreiber, B. Is freezing in the vaccine cold chain an ongoing issue? A literature review. Vaccine 2017, 35, 2127–2133.

(23) Pambudi, N. A.; Sarifudin, A.; Gandidi, I. M.; Romadhon, R. Vaccine cold chain management and cold storage technology to address the challenges of vaccination programs. Energy Reports 2022, 8, 955–972.

(24) Price, W. Advances in Virus Research; Elsevier, 1964; Vol. 10; pp 171–217.

(25) Epand, R. M.; Epand, R. F. Thermal denaturation of influenza virus and its relationship to membrane fusion. Biochemical Journal 2002, 365, 841–848.

(26) Ausar, S. F.; Foubert, T. R.; Hudson, M. H.; Vedvick, T. S.; Middaugh, C. R. Conformational Stability and Disassembly of Norwalk Virus-like Particles. Journal of Biological Chemistry 2006, 281, 19478–19488.

(27) Qiu, X. Heat Induced Capsid Disassembly and DNA Release of Bacteriophage *λ*. PLoS ONE 2012, 7, e39793.

(28) McHugh, C. A.; Tammariello, R. F.; Millard, C. B.; Carra, J. H. Improved stability of a protein vaccine through elimination of a partially unfolded state. Protein Science 2004, 13, 2736–2743.

(29) Dumpa, N.; Goel, K.; Guo, Y.; McFall, H.; Pillai, A. R.; Shukla, A.; Repka, M. A.; Murthy, S. N. Stability of Vaccines. AAPS PharmSciTech 2019, 20, 42.

(30) Kumru, O. S.; Joshi, S. B.; Smith, D. E.; Middaugh, C. R.; Prusik, T.; Volkin, D. B. Vaccine instability in the cold chain: Mechanisms, analysis and formulation strategies. Biologicals 2014, 42, 237–259.

(31) Lloyd, J.; Cheyne, J. The origins of the vaccine cold chain and a glimpse of the future. Vaccine 2017, 35, 2115–2120.

(32) Yu, Y. B.; Briggs, K. T.; Taraban, M. B.; Brinson, R. G.; Marino, J. P. Grand Challenges in Pharmaceutical Research Series: Ridding the Cold Chain for Biologics. Pharm Res 2021, 38, 3–7.

(33) Dadari, I. K.; Zgibor, J. C. How the use of vaccines outside the cold chain or in controlled temperature chain contributes to improving immunization coverage in low-and middle-income countries (LMICs): A scoping review of the literature. J Glob Health 2021, 11, 04004.

(34) Fahrni, M. L.; Ismail, I. A.-N.; Refi, D. M.; Almeman, A.; Yaakob, N. C.; Saman, K. M.; Mansor, N. F.; Noordin, N.; Babar, Z.-U.-D. Management of COVID-19 vaccines cold chain logistics: a scoping review. J of Pharm Policy and Pract 2022, 15, 16.

(35) Hibbs, B. F.; Miller, E.; Shi, J.; Smith, K.; Lewis, P.; Shimabukuro, T. T. Safety of vaccines that have been kept outside of recommended temperatures: Reports to the Vaccine Adverse Event Reporting System (VAERS), 2008–2012. Vaccine 2018, 36, 553–558.

(36) Thielmann, A.; Puth, M.-T.; Weltermann, B. Visual inspection of vaccine storage conditions in general practices: A study of 75 vaccine refrigerators. PLoS ONE 2019, 14, e0225764.

(37) Thielmann, A.; Puth, M.-T.; Weltermann, B. Improving knowledge on vaccine storage management in general practices: Learning effectiveness of an online-based program. Vaccine 2020, 38, 7551–7557.

(38) Gillett, M. B.; Suko, J. R.; Santoso, F. O.; Yancey, P. H. Elevated levels of trimethylamine oxide in muscles of deep-sea gadiform teleosts: A high-pressure adaptation? J. Exp. Zool. 1997, 279, 386–391.

(39) Selvarajan, R.; Sibanda, T.; Tekere, M. Thermophilic bacterial communities inhabiting the microbial mats of “indifferent” and chalybeate (iron-rich) thermal springs: Diversity and biotechnological analysis. MicrobiologyOpen 2018, 7, e00560.

(40) Wen, X.; Wang, S.; Duman, J. G.; Arifin, J. F.; Juwita, V.; Goddard, W. A.; Rios, A.; Liu, F.; Kim, S.-K.; Abrol, R. et al. Antifreeze proteins govern the precipitation of trehalose in a freezing-avoiding insect at low temperature. Proc. Natl. Acad. Sci. U.S.A. 2016, 113, 6683–6688.

(41) Arakawa, T.; Timasheff, S. The stabilization of proteins by osmolytes. Biophysical Journal 1985, 47, 411–414.

(42) Burg, M. B.; Ferraris, J. D. Intracellular Organic Osmolytes: Function and Regulation. Journal of Biological Chemistry 2008, 283, 7309–7313.

(43) Khan, S. H.; Ahmad, N.; Ahmad, F.; Kumar, R. Naturally occurring organic osmolytes: From cell physiology to disease prevention. IUBMB Life 2010, 62, 891–895.

(44) Arakawa, T.; Tsumoto, K.; Kita, Y.; Chang, B.; Ejima, D. Biotechnology applications of amino acids in protein purification and formulations. Amino Acids 2007, 33, 587–605.

(45) Kamerzell, T. J.; Esfandiary, R.; Joshi, S. B.; Middaugh, C. R.; Volkin, D. B. Pro-tein–excipient interactions: Mechanisms and biophysical characterization applied to protein formulation development. Advanced Drug Delivery Reviews 2011, 63, 1118–1159.

(46) Bongioanni, A.; Bueno, M. S.; Mezzano, B. A.; Longhi, M. R.; Garnero, C. Amino acids and its pharmaceutical applications: A mini review. International Journal of Pharmaceutics 2022, 613, 121375.

(47) Jeong, S. H. Analytical methods and formulation factors to enhance protein stability in solution. Arch. Pharm. Res. 2012, 35, 1871–1886.

(48) Ionova, Y.; Wilson, L. Biologic excipients: Importance of clinical awareness of inactive ingredients. PLoS ONE 2020, 15, e0235076.

(49) Kerlin, R. L.; Li, X. Haschek and Rousseaux’s Handbook of Toxicologic Pathology; Elsevier, 2013; pp 725–750.

(50) Kissmann, J.; Ausar, S. F.; Foubert, T. R.; Brock, J.; Switzer, M. H.; Detzi, E. J.; Vedvick, T. S.; Middaugh, C. Physical Stabilization of Norwalk Virus-Like Particles. Journal of Pharmaceutical Sciences 2008, 97, 4208–4218.

(51) Zhang, Y.; Zhang, H.; Ghosh, D. The Stabilizing Excipients in Dry State Therapeutic Phage Formulations. AAPS PharmSciTech 2020, 21, 133.

(52) Pan, L.; Liu, X.; Fan, D.; Qian, Z.; Sun, X.; Wu, P.; Zhong, L. Study of Oncolytic Virus Preservation and Formulation. Pharmaceuticals 2023, 16, 843.

(53) Kim, Y.-S.; Jones, L. S.; Dong, A.; Kendrick, B. S.; Chang, B. S.; Manning, M. C.; Randolph, T. W.; Carpenter, J. F. Effects of sucrose on conformational equilibria and fluctuations within the native-state ensemble of proteins. Protein Sci. 2003, 12, 1252–1261.

(54) Kamerzell, T. J.; Russell Middaugh, C. The Complex Inter-Relationships Between Protein Flexibility and Stability. Journal of Pharmaceutical Sciences 2008, 97, 3494–3517.

(55) Jorgensen, L.; Hostrup, S.; Moeller, E. H.; Grohganz, H. Recent trends in stabilising peptides and proteins in pharmaceutical formulation – considerations in the choice of excipients. Expert Opinion on Drug Delivery 2009, 6, 1219–1230.

(56) Ohtake, S.; Kita, Y.; Arakawa, T. Interactions of formulation excipients with proteins in solution and in the dried state. Advanced Drug Delivery Reviews 2011, 63, 1053–1073, Number: 13.

(57) Barata, T.; Zhang, C.; Dalby, P.; Brocchini, S.; Zloh, M. Identification of Pro-tein–Excipient Interaction Hotspots Using Computational Approaches. IJMS 2016, 17, 853.

(58) Stärtzel, P. Arginine as an Excipient for Protein Freeze-Drying: A Mini Review. Journal of Pharmaceutical Sciences 2018, 107, 960–967.

(59) Castañeda Ruiz, A. J.; Shetab Boushehri, M. A.; Phan, T.; Carle, S.; Garidel, P.; Buske, J.; Lamprecht, A. Alternative Excipients for Protein Stabilization in Protein Therapeutics: Overcoming the Limitations of Polysorbates. Pharmaceutics 2022, 14, 2575.

(60) Santra, S.; Jana, M. Influence of Aqueous Arginine Solution on Regulating Conformational Stability and Hydration Properties of the Secondary Structural Segments of a Protein at Elevated Temperatures: A Molecular Dynamics Study. J. Phys. Chem. B 2022, 126, 1462–1476.

(61) Lošdorfer Božič, A.; Šiber, A.; Podgornik, R. Statistical analysis of sizes and shapes of virus capsids and their resulting elastic properties. J Biol Phys 2013, 39, 215–228.

(62) Ni, T.; Gerard, S.; Zhao, G.; Dent, K.; Ning, J.; Zhou, J.; Shi, J.; Anderson-Daniels, J.; Li, W.; Jang, S. et al. Intrinsic curvature of the HIV-1 CA hexamer underlies capsid topology and interaction with cyclophilin A. Nat Struct Mol Biol 2020, 27, 855–862.

(63) Zhao, G.; Perilla, J. R.; Yufenyuy, E. L.; Meng, X.; Chen, B.; Ning, J.; Ahn, J.; Gronenborn, A. M.; Schulten, K.; Aiken, C. et al. Mature HIV-1 capsid structure by cryo-electron microscopy and all-atom molecular dynamics. Nature 2013, 497, 643–646.

(64) Hadden, J. A.; Perilla, J. R.; Schlicksup, C. J.; Venkatakrishnan, B.; Zlotnick, A.; Schulten, K. All-atom molecular dynamics of the HBV capsid reveals insights into biological function and cryo-EM resolution limits. eLife 2018, 7, e32478.

(65) Quinn, C. M.; Wang, M.; Fritz, M. P.; Runge, B.; Ahn, J.; Xu, C.; Perilla, J. R.; Gronenborn, A. M.; Polenova, T. Dynamic regulation of HIV-1 capsid interaction with the restriction factor TRIM5*α* identified by magic-angle spinning NMR and molecular dynamics simulations. Proc. Natl. Acad. Sci. U.S.A. 2018, 115, 11519–11524.

(66) Ceres, P.; Zlotnick, A. Weak Protein-Protein Interactions Are Sufficient To Drive Assembly of Hepatitis B Virus Capsids. Biochemistry 2002, 41, 11525–11531.

(67) Perilla, J. R.; Hadden, J. A.; Goh, B. C.; Mayne, C. G.; Schulten, K. All-Atom Molecular Dynamics of Virus Capsids as Drug Targets. J. Phys. Chem. Lett. 2016, 7, 1836–1844.

(68) Durrant, J. D.; Kochanek, S. E.; Casalino, L.; Ieong, P. U.; Dommer, A. C.; Amaro, R. E. Mesoscale All-Atom Influenza Virus Simulations Suggest New Substrate Binding Mechanism. ACS Cent. Sci. 2020, 6, 189–196.

(69) Casalino, L.; Seitz, C.; Lederhofer, J.; Tsybovsky, Y.; Wilson, I. A.; Kanekiyo, M.; Amaro, R. E. Breathing and Tilting: Mesoscale Simulations Illuminate Influenza Glycoprotein Vulnerabilities. ACS Cent. Sci. 2022, 8, 1646–1663.

(70) Yu, A.; Pak, A. J.; He, P.; Monje-Galvan, V.; Casalino, L.; Gaieb, Z.; Dommer, A. C.; Amaro, R. E.; Voth, G. A. A multiscale coarse-grained model of the SARS-CoV-2 virion. Biophysical Journal 2021, 120, 1097–1104.

(71) Hagan, M. F.; Zandi, R. Recent advances in coarse-grained modeling of virus assembly. Current Opinion in Virology 2016, 18, 36–43.

(72) Wang, D.; Li, J.; Wang, L.; Cao, Y.; Kang, B.; Meng, X.; Li, S.; Song, C. Toward atomistic models of intact severe acute respiratory syndrome coronavirus 2 via Martini coarse-grained molecular dynamics simulations. Quant. Biol. 2023, 11, 421–433.

(73) Larsson, D. S. D.; Liljas, L.; Van Der Spoel, D. Virus Capsid Dissolution Studied by Microsecond Molecular Dynamics Simulations. PLoS Comput Biol 2012, 8, e1002502.

(74) Roberts, J. A.; Kuiper, M. J.; Thorley, B. R.; Smooker, P. M.; Hung, A. Investigation of a predicted N-terminal amphipathic *α*-helix using atomistic molecular dynamics simulation of a complete prototype poliovirus virion. Journal of Molecular Graphics and Modelling 2012, 38, 165–173.

(75) Arcangeli, C.; Circelli, P.; Donini, M.; Aljabali, A. A.; Benvenuto, E.; Lomonossoff, G. P.; Marusic, C. Structure-based design and experimental engineering of a plant virus nanoparticle for the presentation of immunogenic epitopes and as a drug carrier. Journal of Biomolecular Structure and Dynamics 2014, 32, 630–647.

(76) Perilla, J. R.; Schulten, K. Physical properties of the HIV-1 capsid from all-atom molecular dynamics simulations. Nat Commun 2017, 8, 15959.

(77) Pohjolainen, E.; Malola, S.; Groenhof, G.; Häkkinen, H. Exploring Strategies for Labeling Viruses with Gold Nanoclusters through Non-equilibrium Molecular Dynamics Simulations. Bioconjugate Chem. 2017, 28, 2327–2339.

(78) Casalino, L.; Dommer, A.; Gaieb, Z.; Barros, E. P.; Sztain, T.; Ahn, S.-H.; Trifan, A.; Brace, A.; Bogetti, A.; Ma, H. et al. AI-Driven Multiscale Simulations Illuminate Mechanisms of SARS-CoV-2 Spike Dynamics. 2020.

(79) Li, C.; Chen, W.; Lin, X.; Zhang, S.; Wang, Y.; He, X.; Ren, Y. Molecular dynamics study on the stability of foot-and-mouth disease virus particle in salt solution. Molecular Simulation 2021, 47, 1104–1111.

(80) Dommer, A.; Casalino, L.; Kearns, F.; Rosenfeld, M.; Wauer, N.; Ahn, S.-H.; Russo, J.; Oliveira, S.; Morris, C.; Bogetti, A. et al. #COVIDisAirborne: AI-enabled multi-scale computational microscopy of delta SARS-CoV-2 in a respiratory aerosol. The International Journal of High Performance Computing Applications 2023, 37, 28–44.

(81) Shrivastav, G.; Borkotoky, S.; Dey, D.; Singh, B.; Malhotra, N.; Azad, K.; Jayaram, B.; Agarwal, M.; Banerjee, M. Structure and energetics guide dynamic behaviour in a T = 3 icosahedral virus capsid. Biophysical Chemistry 2024, 305, 107152.

(82) Ghaemi, Z.; Gruebele, M.; Tajkhorshid, E. Molecular mechanism of capsid disassembly in hepatitis B virus. Proc. Natl. Acad. Sci. U.S.A. 2021, 118, e2102530118.

(83) Humphrey, W.; Dalke, A.; Schulten, K. VMD: Visual molecular dynamics. Journal of Molecular Graphics 1996, 14, 33–38.

(84) Simpson, A. A.; Hébert, B.; Sullivan, G. M.; Parrish, C. R.; Zádori, Z.; Tijssen, P.; Rossmann, M. G. The structure of porcine parvovirus: comparison with related viruses. Journal of Molecular Biology 2002, 315, 1189–1198.

(85) Roos, W. H.; Ivanovska, I. L.; Evilevitch, A.; Wuite, G. J. L. Viral capsids: Mechanical characteristics, genome packaging and delivery mechanisms. Cell. Mol. Life Sci. 2007, 64, 1484.

(86) Selivanovitch, E.; LaFrance, B.; Douglas, T. Molecular exclusion limits for diffusion across a porous capsid. Nat Commun 2021, 12, 2903.

(87) Inoue, H.; Timasheff, S. N. Preferential and absolute interactions of solvent components with proteins in mixed solvent systems. Biopolymers 1972, 11, 737–743.

(88) Record, M.; Anderson, C. Interpretation of preferential interaction coefficients of nonelectrolytes and of electrolyte ions in terms of a two-domain model. Biophysical Journal 1995, 68, 786–794.

(89) Shukla, D.; Shinde, C.; Trout, B. L. Molecular Computations of Preferential Interaction Coefficients of Proteins. J. Phys. Chem. B 2009, 113, 12546–12554.

(90) Vagenende, V.; Yap, M. G. S.; Trout, B. L. Molecular Anatomy of Preferential Interaction Coefficients by Elucidating Protein Solvation in Mixed Solvents: Methodology and Application for Lysozyme in Aqueous Glycerol. J. Phys. Chem. B 2009, 113, 11743–11753.

(91) Van Der Spoel, D.; Lindahl, E.; Hess, B.; Groenhof, G.; Mark, A. E.; Berendsen, H. J. C. GROMACS: Fast, flexible, and free. J. Comput. Chem. 2005, 26, 1701–1718.

(92) Abraham, M. J.; Murtola, T.; Schulz, R.; Páll, S.; Smith, J. C.; Hess, B.; Lindahl, E. GROMACS: High performance molecular simulations through multi-level parallelism from laptops to supercomputers. SoftwareX 2015, 1–2, 19–25.

(93) Strande, S.; Cai, H.; Tatineni, M.; Pfeiffer, W.; Irving, C.; Majumdar, A.; Mishin, D.; Sinkovits, R.; Norman, M.; Wolter, N. et al. Expanse: Computing without Boundaries: Architecture, Deployment, and Early Operations Experiences of a Supercomputer Designed for the Rapid Evolution in Science and Engineering. Practice and Experience in Advanced Research Computing. Boston MA USA, 2021; pp 1–4.

(94) Wardi, Y. A stochastic steepest-descent algorithm. J Optim Theory Appl 1988, 59, 307–323.

(95) Bussi, G.; Donadio, D.; Parrinello, M. Canonical sampling through velocity rescaling. J. Chem. Phys. 2007, 126, 014101.

(96) Evans, D. J.; Holian, B. L. The Nose–Hoover thermostat. J. Chem. Phys. 1985, 83, 4069–4074.

(97) Darden, T.; York, D.; Pedersen, L. Particle mesh Ewald: An *N* •log(*N*) method for Ewald sums in large systems. The Journal of Chemical Physics 1993, 98, 10089–10092.

(98) Yeh, I.-C.; Berkowitz, M. L. Ewald summation for systems with slab geometry. The Journal of Chemical Physics 1999, 111, 3155–3162.

(99) Lorentz, H. A. Ueber die Anwendung des Satzes vom virial in der Kinetischen Theorie der Gase. Annalen der Physik 1881, 248, 127–136.

(100) Berthelot, D. Sur le mélange des gaz. Compt. Rendus 1898, 126.

(101) Brooks, B. R.; Brooks, C. L.; Mackerell, A. D.; Nilsson, L.; Petrella, R. J.; Roux, B.; Won, Y.; Archontis, G.; Bartels, C.; Boresch, S. et al. CHARMM: The biomolecular simulation program. J. Comput. Chem. 2009, 30, 1545–1614.

(102) Huang, J.; MacKerell, A. D. CHARMM36 all-atom additive protein force field: Validation based on comparison to NMR data. J. Comput. Chem. 2013, 34, 2135–2145.

(103) Cloutier, T.; Sudrik, C.; Sathish, H. A.; Trout, B. L. Kirkwood–Buff-derived alcohol parameters for aqueous carbohydrates and their application to preferential interaction coefficient calculations of proteins. The Journal of Physical Chemistry B 2018, 122, 9350–9360.

(104) Jorgensen, W. L.; Chandrasekhar, J.; Madura, J. D.; Impey, R. W.; Klein, M. L. Comparison of simple potential functions for simulating liquid water. The Journal of Chemical Physics 1983, 79, 926–935.

(105) Olsson, M. H. M.; Søndergaard, C. R.; Rostkowski, M.; Jensen, J. H. PROPKA3: Consistent Treatment of Internal and Surface Residues in Empirical p *K* _a_ Predictions. J. Chem. Theory Comput. 2011, 7, 525–537.

(106) McCammon, J. A. Protein dynamics. Rep. Prog. Phys. 1984, 47, 1–46.

(107) Grewal, R.; Roy, S. Modeling proteins as residue interaction networks. PPL 2015, 22, 923–933.

(108) Franke, L.; Peter, C. Visualizing the Residue Interaction Landscape of Proteins by Temporal Network Embedding. J. Chem. Theory Comput. 2023, 19, 2985–2995.

(109) Dasetty, S.; Zajac, J. W. P.; Sarupria, S. Exploitation of active site flexibility-low temperature activity relation for engineering broad range temperature active enzymes. Mol. Syst. Des. Eng. 2023, 8, 1355–1370.

(110) Yehorova, D.; Di Geronimo, B.; Robinson, M.; Kasson, P. M.; Kamerlin, S. C. Using residue interaction networks to understand protein function and evolution and to engineer new proteins. Current Opinion in Structural Biology 2024, 89, 102922.

(111) Freeman, L. C. Centrality in social networks conceptual clarification. Social Networks 1978, 1, 215–239.

(112) Brandes, U. A faster algorithm for betweenness centrality. The Journal of Mathematical Sociology 2001, 25, 163–177.

(113) Brandes, U. On variants of shortest-path betweenness centrality and their generic computation. Social Networks 2008, 30, 136–145.

(114) Wadell, H. Volume, Shape, and Roundness of Quartz Particles. The Journal of Geology 1935, 43, 250–280.

(115) Trimesh [Computer software]. (2019). Retrieved from https://github.com/mikedh/trimesh.

(116) Mi, X.; Blocher McTigue, W. C.; Joshi, P. U.; Bunker, M. K.; Heldt, C. L.; Perry, S. L. Thermostabilization of viruses *via* complex coacervation. Biomater. Sci. 2020, 8, 7082–7092.

(117) Joshi, P. U.; Decker, C.; Zeng, X.; Sathyavageeswaran, A.; Perry, S. L.; Heldt, C. L. Design Rules for the Sequestration of Viruses into Polypeptide Complex Coacervates. Biomacromolecules 2024, 25, 741–753.

(118) Streck, A. F.; Truyen, U. Porcine Parvovirus. Current Issues in Molecular Biology 2020, 33–46.

(119) Gencoglu, M. F.; Pearson, E.; Heldt, C. L. Porcine parvovirus flocculation and removal in the presence of osmolytes. Journal of Biotechnology 2014, 186, 83–90.

(120) Tafur, M. F.; Vijayaragavan, K. S.; Heldt, C. L. Reduction of porcine parvovirus infectivity in the presence of protecting osmolytes. Antiviral Research 2013, 99, 27–33.

(121) Agarwal, H.; Wang, X.; Kulkarni, N. R.; Tao, S.; Demers, C. Application of machine learning in ensuring viral safety of biotherapeutics: Case study demonstrating prediction and optimization of viral clearance performance of anion exchange chromatography. Current Research in Biotechnology 2023, 6, 100140.

(122) Nowak, T.; Popp, B.; Roth, N. J. Choice of parvovirus model for validation studies influences the interpretation of the effectiveness of a virus filtration step. Biologicals 2019, 60, 85–92.

(123) Miesegaes, G.; Lute, S.; Brorson, K. Analysis of viral clearance unit operations for monoclonal antibodies. Biotech & Bioengineering 2010, 106, 238–246.

(124) Rueda, P.; Fominaya, J.; Langeveld, J. P.; Bruschke, C.; Vela, C.; Casal, J. Effect of different baculovirus inactivation procedures on the integrity and immunogenicity of porcine parvovirus-like particles. Vaccine 2000, 19, 726–734.

(125) Marrink, S. J.; Tieleman, D. P. Perspective on the Martini model. Chem. Soc. Rev. 2013, 42, 6801.

(126) Marrink, S. J.; Monticelli, L.; Melo, M. N.; Alessandri, R.; Tieleman, D. P.; Souza, P. C. T. Two decades of Martini: Better beads, broader scope. WIREs Comput Mol Sci 2023, 13, e1620.

(127) Van Vlijmen, H. W.; Karplus, M. Normal Mode Calculations of Icosahedral Viruses with Full Dihedral Flexibility by Use of Molecular Symmetry. Journal of Molecular Biology 2005, 350, 528–542.

(128) Pathak, A. K.; Bandyopadhyay, T. Heat-induced transitions of an empty minute virus of mice capsid in explicit water: all-atom MD simulation. Journal of Biomolecular Structure and Dynamics 2022, 40, 11900–11913.

(129) Szoverfi, J.; Fejer, S. N. Dynamic stability of salt stable cowpea chlorotic mottle virus capsid protein dimers and pentamers of dimers. Sci Rep 2022, 12, 14251.

(130) Cieplak, M.; Robbins, M. O. Nanoindentation of 35 Virus Capsids in a Molecular Model: Relating Mechanical Properties to Structure. PLoS ONE 2013, 8, e63640.

(131) Boyd, K. J.; Bansal, P.; Feng, J.; May, E. R. Stability of Norwalk Virus Capsid Protein Interfaces Evaluated by in Silico Nanoindentation. Front. Bioeng. Biotechnol. 2015, 3.

(132) Gibbons, M. M.; Klug, W. S. Influence of Nonuniform Geometry on Nanoindentation of Viral Capsids. Biophysical Journal 2008, 95, 3640–3649.

(133) Michel, J. P.; Ivanovska, I. L.; Gibbons, M. M.; Klug, W. S.; Knobler, C. M.; Wuite, G. J. L.; Schmidt, C. F. Nanoindentation studies of full and empty viral capsids and the effects of capsid protein mutations on elasticity and strength. Proc. Natl. Acad. Sci. U.S.A. 2006, 103, 6184–6189.

(134) Mi, X.; Heldt, C. L. Single-Particle Chemical Force Microscopy to Characterize Virus Surface Chemistry. BioTechniques 2020, 69, 363–370.

(135) Mi, X.; Bromley, E. K.; Joshi, P. U.; Long, F.; Heldt, C. L. Virus Isoelectric Point Determination Using Single-Particle Chemical Force Microscopy. Langmuir 2020, 36, 370–378.

(136) Zlotnick, A.; Aldrich, R.; Johnson, J. M.; Ceres, P.; Young, M. J. Mechanism of Capsid Assembly for an Icosahedral Plant Virus. Virology 2000, 277, 450–456, Publisher: Elsevier BV.

(137) Mateu, M. G. Assembly, stability and dynamics of virus capsids. Archives of Biochemistry and Biophysics 2013, 531, 65–79.

(138) Du, C.; Cleary, S. P.; Kostelic, M. M.; Jones, B. J.; Kafader, J. O.; Wysocki, V. H. Combining Surface-Induced Dissociation and Charge Detection Mass Spectrometry to Reveal the Native Topology of Heterogeneous Protein Complexes. Anal. Chem. 2023, 95, 13889–13896.

(139) Sherman, M. B.; Smith, H. Q.; Smith, T. J. The Dynamic Life of Virus Capsids. Viruses 2020, 12, 618.

(140) Dong, Y.; Liu, Y.; Jiang, W.; Smith, T. J.; Xu, Z.; Rossmann, M. G. Antibody-induced uncoating of human rhinovirus B14. Proc. Natl. Acad. Sci. U.S.A. 2017, 114, 8017–8022.

(141) Bostina, M.; Levy, H.; Filman, D. J.; Hogle, J. M. Poliovirus RNA Is Released from the Capsid near a Twofold Symmetry Axis. J Virol 2011, 85, 776–783.

(142) Rayaprolu, V.; Kruse, S.; Kant, R.; Venkatakrishnan, B.; Movahed, N.; Brooke, D.; Lins, B.; Bennett, A.; Potter, T.; McKenna, R. et al. Comparative Analysis of Adeno-Associated Virus Capsid Stability and Dynamics. J Virol 2013, 87, 13150–13160.

(143) Reguera, J.; Carreira, A.; Riolobos, L.; Almendral, J. M.; Mateu, M. G. Role of interfacial amino acid residues in assembly, stability, and conformation of a spherical virus capsid. Proc. Natl. Acad. Sci. U.S.A. 2004, 101, 2724–2729.

(144) Zhang, X.; Jiang, L.; Weng, G.; Shen, C.; Zhang, O.; Liu, M.; Zhang, C.; Gu, S.; Wang, J.; Wang, X. et al. HawkDock version 2: an updated web server to predict and analyze the structures of protein–protein complexes. Nucleic Acids Research 2025, 53, W306–W315.

(145) Sun, Y.; Shen, Z.; Zhang, C.; Yi, Y.; Zhu, K.; Xu, F.; Kong, W. Development of a Stable Liquid Formulation for Live Attenuated Influenza Vaccine. Journal of Pharmaceutical Sciences 2019, 108, 2315–2322.

(146) Ohtake, S.; Martin, R.; Saxena, A.; Pham, B.; Chiueh, G.; Osorio, M.; Kopecko, D.; Xu, D.; Lechuga-Ballesteros, D.; Truong-Le, V. Room temperature stabilization of oral, live attenuated Salmonella enterica serovar Typhi-vectored vaccines. Vaccine 2011, 29, 2761–2771.

(147) Tonnis, W.; Amorij, J.-P.; Vreeman, M.; Frijlink, H.; Kersten, G.; Hinrichs, W. Improved storage stability and immunogenicity of hepatitis B vaccine after spray-freeze drying in presence of sugars. European Journal of Pharmaceutical Sciences 2014, 55, 36–45.

(148) Pachauri, R. Stability of live attenuated classical swine fever cell culture vaccine virus in liquid form for developing an oral vaccine. 2020,

(149) Maginnis, M. S. Virus–Receptor Interactions: The Key to Cellular Invasion. Journal of Molecular Biology 2018, 430, 2590–2611.

(150) Su, R.; Zeng, J.; Marcink, T. C.; Porotto, M.; Moscona, A.; O’Shaughnessy, B. Host Cell Membrane Capture by the SARS-CoV-2 Spike Protein Fusion Intermediate. ACS Cent. Sci. 2023, 9, 1213–1228.

(151) Pipatpadungsin, N.; Chao, K.; Rouse, S. L. Coarse-Grained Simulations of Adeno-Associated Virus and Its Receptor Reveal Influences on Membrane Lipid Organization and Curvature. J. Phys. Chem. B 2024, 128, 10139–10153.

(152) Peccati, F.; Jiménez-Osés, G. Enthalpy–Entropy Compensation in Biomolecular Recognition: A Computational Perspective. ACS Omega 2021, 6, 11122–11130.

(153) Dong, J.; Davis, A. P. Molecular Recognition Mediated by Hydrogen Bonding in Aqueous Media. Angewandte Chemie 2021, 133, 8113–8126.

(154) Zhu, Y.; Tang, M.; Zhang, H.; Rahman, F.-U.; Ballester, P.; Rebek, J.; Hunter, C. A.; Yu, Y. Water and the Cation Interaction. J. Am. Chem. Soc. 2021, 143, 12397–12403.

(155) Cole, A. G. Modulators of HBV capsid assembly as an approach to treating hepatitis B virus infection. Current Opinion in Pharmacology 2016, 30, 131–137.

(156) Liu, K.; Du, S.; Deng, W.; Qu, Z.; Hu, X. Advances in the computational development of hepatitis B virus capsid assembly modulators. Drug Discovery Today 2025, 30, 104458.

(157) Pavlova, A.; Bassit, L.; Cox, B. D.; Korablyov, M.; Chipot, C.; Patel, D.; Lynch, D. L.; Amblard, F.; Schinazi, R. F.; Gumbart, J. C. The Mechanism of Action of Hepatitis B Virus Capsid Assembly Modulators Can Be Predicted from Binding to Early Assembly Intermediates. J. Med. Chem. 2022, 65, 4854–4864.

(158) Pavlova, A.; Fan, Z.; Lynch, D. L.; Gumbart, J. C. Machine Learning of Molecular Dynamics Simulations Provides Insights into the Modulation of Viral Capsid Assembly. J. Chem. Inf. Model. 2025, 65, 4844–4853.

(159) Askari, F. S.; Mohebbi, A. The dynamic states of hepatitis B virus capsid monomers under the impact of different class of capsid-assembly modulators. Sci Rep 2025, 15, 32731.

(160) Sevvana, M.; Klose, T.; Rossmann, M. G. Encyclopedia of Virology; Elsevier, 2021; pp 257–277.

(161) Yoneda, S.; Kitazawa, M.; Umeyama, H. Molecular dynamics simulation of a rhinovirus capsid under rotational symmetry boundary conditions. J. Comput. Chem. 1996, 17, 191–203.

(162) Roy, A.; Post, C. B. Microscopic Symmetry Imposed by Rotational Symmetry Boundary Conditions in Molecular Dynamics Simulation. J. Chem. Theory Comput. 2011, 7, 3346–3353.

(163) Roy, A.; Post, C. B. Long-distance correlations of rhinovirus capsid dynamics contribute to uncoating and antiviral activity. Proc. Natl. Acad. Sci. U.S.A. 2012, 109, 5271–5276.

(164) Li, J.; Chang, Y.; Yang, S.; Zhou, G.; Wei, Y. Formulation enhanced the stability of Foot-and-mouth virus and prolonged vaccine storage. Virol J 2022, 19, 207.

(165) Wiggan, O.; Livengood, J. A.; Silengo, S. J.; Kinney, R. M.; Osorio, J. E.; Huang, C. Y.-H.; Stinchcomb, D. T. Novel formulations enhance the thermal stability of live-attenuated flavivirus vaccines. Vaccine 2011, 29, 7456–7462.

(166) Schlehuber, L. D.; McFadyen, I. J.; Shu, Y.; Carignan, J.; Duprex, W. P.; Forsyth, W. R.; Ho, J. H.; Kitsos, C. M.; Lee, G. Y.; Levinson, D. A. et al. Towards ambient temperature-stable vaccines: The identification of thermally stabilizing liquid formulations for measles virus using an innovative high-throughput infectivity assay. Vaccine 2011, 29, 5031–5039.

(167) Ohtake, S.; Arakawa, T.; Koyama, A. H. Arginine as a Synergistic Virucidal Agent. Molecules 2010, 15, 1408–1424.

(168) Meingast, C.; Heldt, C. L. Arginine-enveloped virus inactivation and potential mechanisms. Biotechnology Progress 2020, 36, e2931.

(169) Meingast, C. L.; Joshi, P. U.; Turpeinen, D. G.; Xu, X.; Holstein, M.; Feroz, H.; Ranjan, S.; Ghose, S.; Li, Z. J.; Heldt, C. L. Physiochemical properties of enveloped viruses and arginine dictate inactivation. Biotechnology Journal 2021, 16, 2000342.

(170) Zajac, J. W. P.; Muralikrishnan, P.; Tohidian, I.; Zeng, X.; Heldt, C. L.; Perry, S. L.; Sarupria, S. Flipping out: role of arginine in hydrophobic interactions and biological formulation design. Chem. Sci. 2025, 16, 6780–6792.

(171) Barbour, E.; Abdelnour, A.; Jirjis, F.; Faroon, O.; Farran, M. Evaluation of 12 stabilizers in a developed attenuated Salmonella Enteritidis vaccine. Vaccine 2002, 20, 2249–2253.

(172) White, J. A.; Estrada, M.; Flood, E. A.; Mahmood, K.; Dhere, R.; Chen, D. Development of a stable liquid formulation of live attenuated influenza vaccine. Vaccine 2016, 34, 3676–3683.

