## Supporting Information for "Cracking the Capsid Code: A Computationally-Feasible Approach for Investigating Virus-Excipient Interactions in Biologics Design"

### Contents


**1 Capsid Surface Construction** **S-3**

**2 Equilibration and Convergence** **S-5**

**3 Water Structure and Dynamics** **S-9**

3.1 Three-Body Angle Distributions . . . . . . . . . . . . . . . . . . . . . . . . . . . . S-9

3.2 Water Reorientation Dynamics . . . . . . . . . . . . . . . . . . . . . . . . . . . . S-9

3.3 Justification of $h_{box}$ . . . . . . . . . . . . . . . . . . . . . . . . . . . . . . . . . . . S-10

**4 Pulling Simulations** **S-13**

**5 Performance** **S-18**

**6 Capsid Structure and Dynamics** **S-19**

**7 Experimental Details** **S-22**

**References** **S-24**

### 1 Capsid Surface Construction

The first stage of the structure construction module involves a symmetry-based alignment of an input capsid structure (Fig. 1b). In this stage, capsid Cartesian coordinates, $\mathbf{r}_{\text{Capsid}}$, are transformed by a translation vector, $\mathbf{T}$, such that the center-of-geometry of reference atoms $\mathbf{r}_{\text{COG},Ref}$ is at the origin:

$$\mathbf{r}_{\text{COG,Ref}} = \frac{1}{3}(\mathbf{r}_1 + \mathbf{r}_2 + \mathbf{r}_3) \tag{S1}$$

$$\mathbf{T}_{\mathbf{r}} = \mathbf{r}_{\text{Capsid}} - \mathbf{r}_{\text{COG,Ref}} \tag{S2}$$

where $\mathbf{r}_i$ is the vector of each point from the origin. Following translation, the normal, $\mathbf{n}_{\mathbf{ref}}$, to the plane connecting the three reference atoms is computed as:

$$\mathbf{n}_{\mathbf{ref}} = \frac{(\mathbf{r}_2 - \mathbf{r}_1) \times (\mathbf{r}_3 - \mathbf{r}_1)}{|(\mathbf{r}_2 - \mathbf{r}_1) \times (\mathbf{r}_3 - \mathbf{r}_1)|} \tag{S3}$$

which is used to define a rotation matrix, $\mathbf{R}$, applied to capsid coordinates such that the normal is aligned with the $z$-axis, $\mathbf{e}_z$:

$$\theta = \cos^{-1}(\mathbf{n}_{\mathbf{ref}} \cdot \mathbf{e}_z) \tag{S4}$$

$$R = I + (\sin\theta)[\mathbf{K}] + (1 - \cos\theta)[\mathbf{K}]^2 \tag{S5}$$

where $I$ is the $3 \times 3$ identity matrix and $\mathbf{K}$ is the skew-symmetric matrix of $\mathbf{n}_{\mathbf{ref}}$.

Following alignment, the capsid structure is converted into a surface model (Fig. 1c). In this stage, (i) the capsid is shifted such that its center-of-geometry is at the origin:

$$\mathbf{T}_{\mathbf{c}} = \mathbf{r}_{\text{Capsid}} - \mathbf{r}_{\text{COG,Capsid}} \tag{S6}$$

Fig. S1 shows a demonstration of the CapSACIN surface abstraction module to six distinct capsid architectures: Sulfolobus Turreted Icosahedral Virus (PDB: 3J31), Infectious Bursal Virus (PDB: 2GSY), Maize Chlorotic Mottle Virus (PDB: 3JB8), Foot-and-Mouth-

Disease Virus (PDB: 8HBG), Human Papillomavirus Type 16 (PDB: 3J6R), and Saccharomyces Cerevisiae Virus L-BCLa (PDB: 7QWZ).

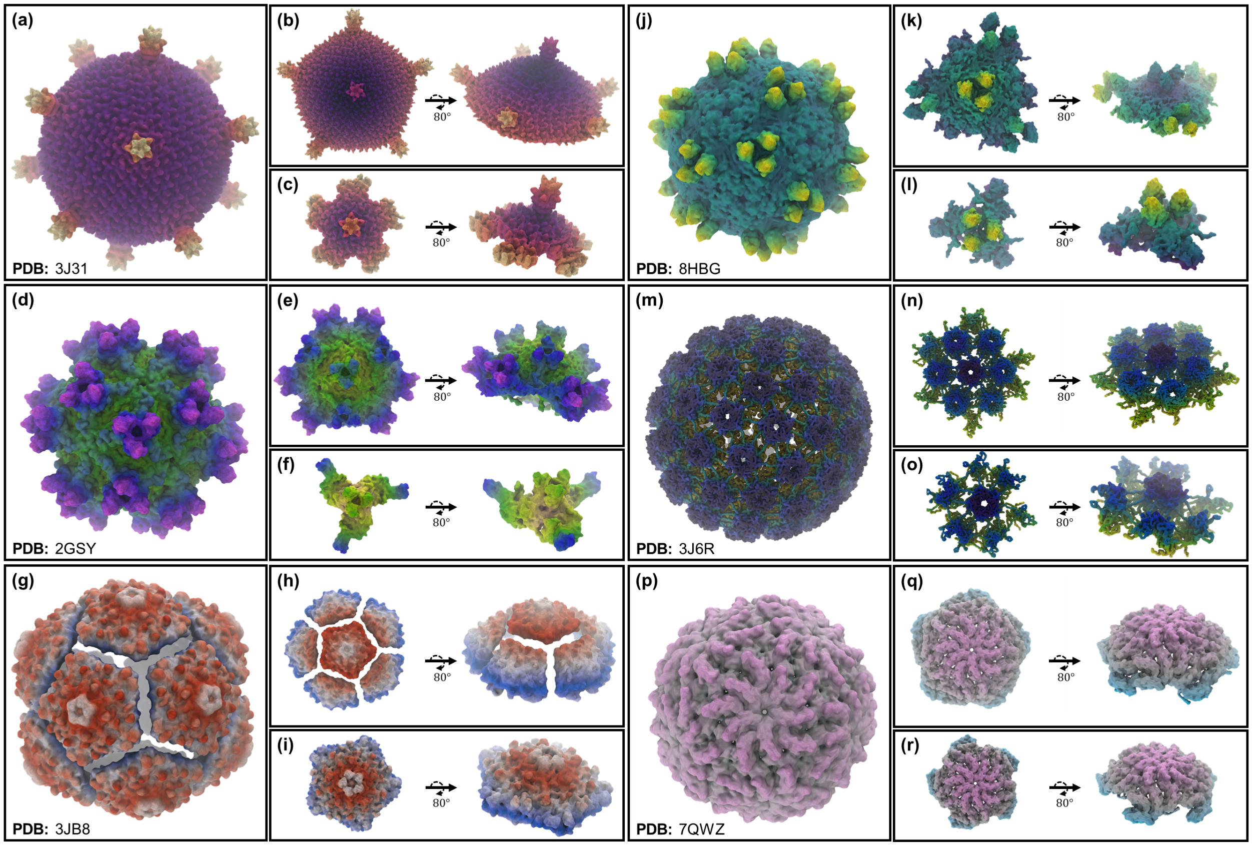


**Figure S1:** Capsid surface model generation for six unique capsids available from the Protein Data Bank. (a) Sulfolobus Turreted Icosahedral Virus (PDB: 3J31). (d) Infectious Bursal Virus (PDB: 2GSY). (g) Maize Chlorotic Mottle Virus (PDB: 3JB8). (j) Foot-and-Mouth-Disease Virus (PDB: 8HBG). (m) Human Papillomavirus Type 16 (PDB: 3J6R). (p) Saccharomyces Cerevisiae Virus L-BCLa (PDB: 7QWZ). (b,e,h,k,n,q) Moderately truncated virus surface models (between 40-70% system size reduction). (c,f,i,l,o,r) Significantly truncated virus surface models (between 70-90% system size reduction).

### 2 Equilibration and Convergence

Excipient positions are initialized as a 0.1 nm thick slab at the top of the simulation box (Fig. S2a). This slab is used as the reference position for the flat-bottom restraint described in Section 2.1.3. After equilibration and during production runs, no excipient molecules are found to interact with the restrained portion of the capsid surface (Fig. S2b,c). The capsid position restraining scheme used during equilibration is shown in Fig. S2a.

**Table S1:** PPV systems under study. Numbers shown include the number of capsid protein monomers ($N_{CP}$), water molecules ($N_{WAT}$), excipient molecules ($N_{Exc}$), sodium ions ($N_{Na^+}$), and chloride ions ($N_{Cl^-}$).

| **System** | $N_{CP}$ | $N_{WAT}$ | $N_{Exc}$ | $N_{Na^+}$ | $N_{Cl^-}$ |
|---|---|---|---|---|---|
| PPV Assembled Capsid | 60 | 877,837 | 0 | 3,338 | 2,858 |
| PPV 5-fold Surface | 15 | 205,697 | 0 | 799 | 679 |
| PPV 3-fold Surface | 9 | 186,162 | 0 | 653 | 581 |
| PPV 2-fold Surface | 12 | 196,683 | 0 | 728 | 632 |
| PPV 5-fold Surface / 0.1 M ARG | 15 | 201,665 | 396 | 679 | 955 |
| PPV 5-fold Surface / 0.1 M TRE | 15 | 199,204 | 396 | 799 | 679 |
| PPV 5-fold Surface / 0.1 M GLY | 15 | 203,971 | 396 | 799 | 679 |
| PPV 5-fold Surface / 0.1 M SOR | 15 | 201,909 | 396 | 799 | 679 |
| PPV 5-fold Surface / 0.1 M GLU | 15 | 202,508 | 396 | 1195 | 679 |

Excipient convergence was measured *via* preferential interaction coefficients, $\Gamma_{XV}$, calculated according to the two-domain formula[S1–S3] given by:

$$\Gamma_{XV} = \left\langle N_X^{\text{local}} - \left( \frac{N_X^{\text{bulk}}}{N_W^{\text{bulk}}} \right) N_W^{\text{local}} \right\rangle \tag{S7}$$

where $V$ denotes the viral surface, $X$ represents an excipient species, and $W$ indicates water. $N$ represents the number of molecules of a given species in the local or bulk domain of the viral surface. $N$ represents the number of molecules of a given species, while angular brackets denote an ensemble average. The local and bulk domain is separated by a cutoff distance

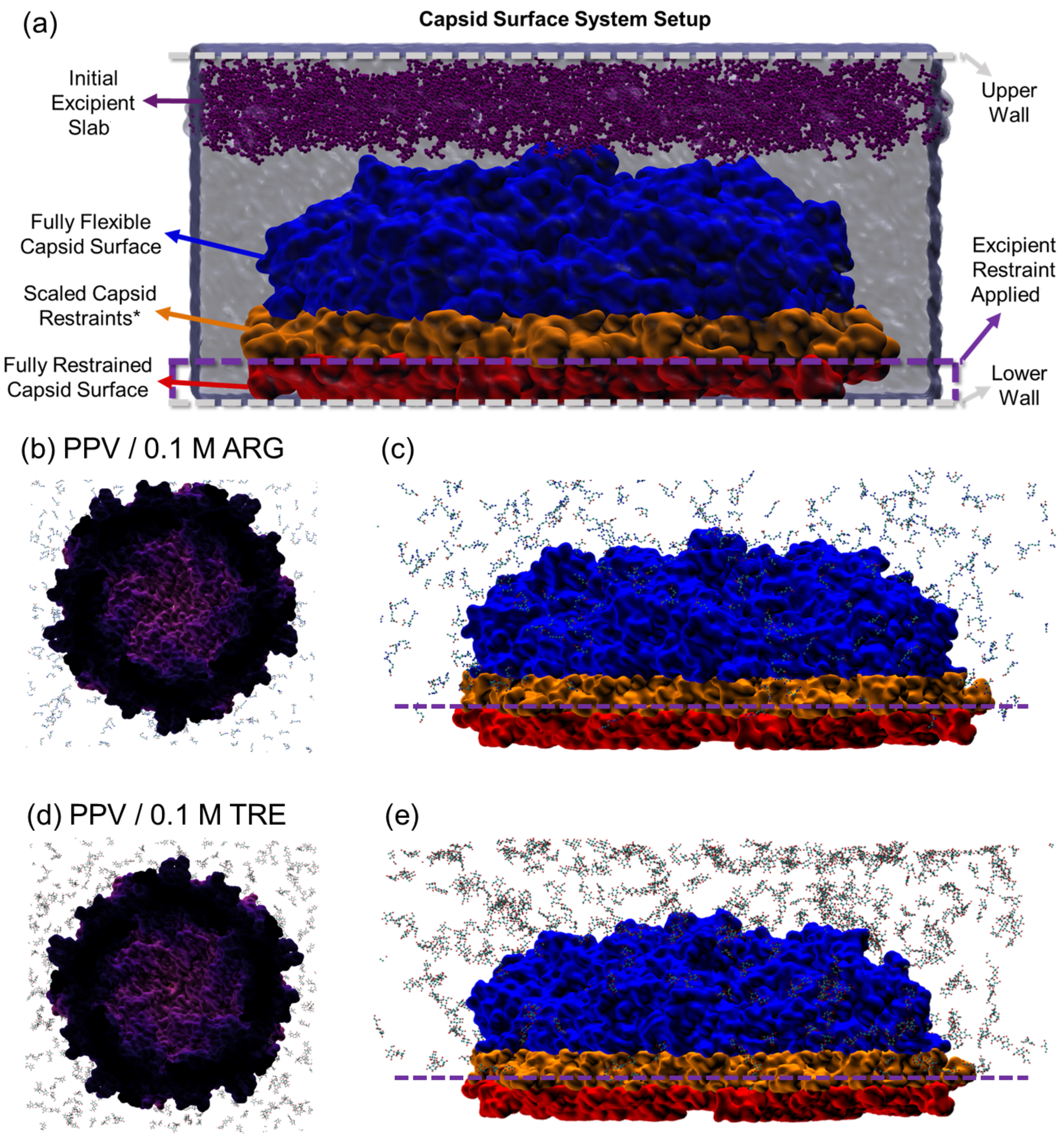


**Figure S2:** Excipient initialization and position restraint scheme for capsid surface construction. (a) Overview of the position restraining scheme, including the initial excipient 1.0 nm thick slab at the top of the simulation box. Simulation snapshots are shown for the PPV 5-fold surface in (b) 0.1 M arginine (ARG) and (c) 0.1 M trehalose (TRE) solutions, showing a depletion of excipient molecules from the restrained regions.

$R_{cut}$ from the capsid surface. $\Gamma_{XV}$ gives a measure of the relative accumulation or depletion of an additive in the local domain of the virus; $\Gamma_{XV} > 0$ indicates relative accumulation (preferential interaction) and $\Gamma_{XV} < 0$ indicates relative depletion (preferential exclusion).

We observed that, at all three ROIs, ARG preferentially interacts with the virus surface while TRE is preferentially excluded (Fig. S3). More importantly, our values of $\Gamma_{XV}$ are observed to converge to constant values both with increasing $R_{cut}$ distances and increased simulation time. In Fig. S5c, we show $\Gamma_{XV}$ fluctuates around a mean value for all excipient/5-fold surface simulations ($R_{cut} = 0.7$ nm), indicating equilibrium of the solvent distribution. Fig. S5a,b demonstrate structural equilibrium of the capsid surface for these same systems.

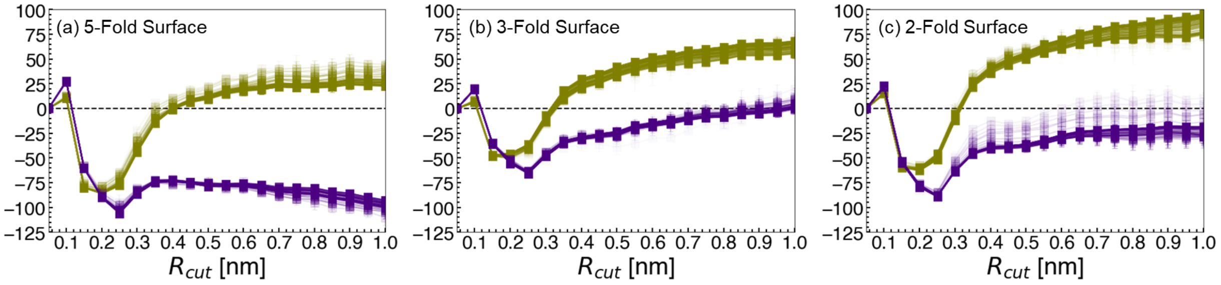


**Figure S3:** $\Gamma_{XV}$ values for excipient-virus preferential interactions. (a) 5-fold, (b) 3-fold, and (c) 2-fold surfaces. $\Gamma_{XV}$ values for ARG are colored in olive, while TRE is shown in indigo. Increased shading indicates increased simulation time, ranging from 0-50 ns.

We assess conformational equilibration using a local measure and global measure. The local measure uses the root-mean-square deviation of $C_\alpha$ atoms, as:

$$RMSD_{C_\alpha}(t) = \left[\frac{1}{N}\sum_{i=1}^{N}(\mathbf{r}_i(t) - \mathbf{r}_i(t_0))^2\right]^{\frac{1}{2}} \tag{S8}$$

In the fully assembled PPV capsid, all monomers individually have $RMSD_{C_\alpha}$ fluctuations between 0.125-0.24 nm (Fig. S4a). This includes 5-fold ROI monomers that are later used to compare with surface model data. In the 5-fold surface model, 5-fold ROI monomers have $RMSD_{C_\alpha}$ fluctuations within a range of the same width (0.1-0.215 nm), while peripheral monomers are subject to much more significant conformational deviations (Fig. S4b).

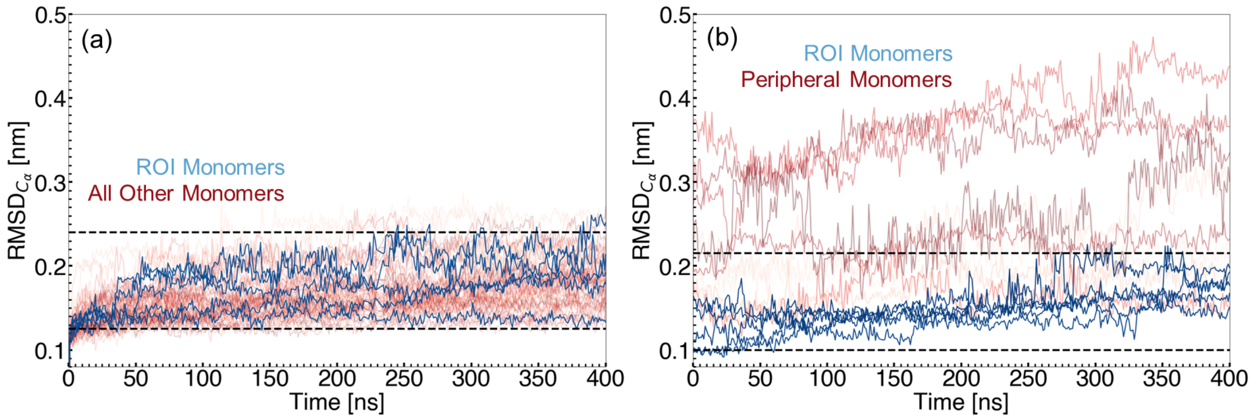


**Figure S4:** RMSD of $C_\alpha$ atoms in PPV monomers. (a) Fully assembled PPV capsid. (b) 5-fold surface model. RMSD of monomers extracted from the 5-fold ROI are shown in blue, while other capsid protein monomers are colored in red. Dashed horizontal lines are used to approximate the range in which $RMSD_{C_\alpha}$ varies in the assembled PPV capsid.

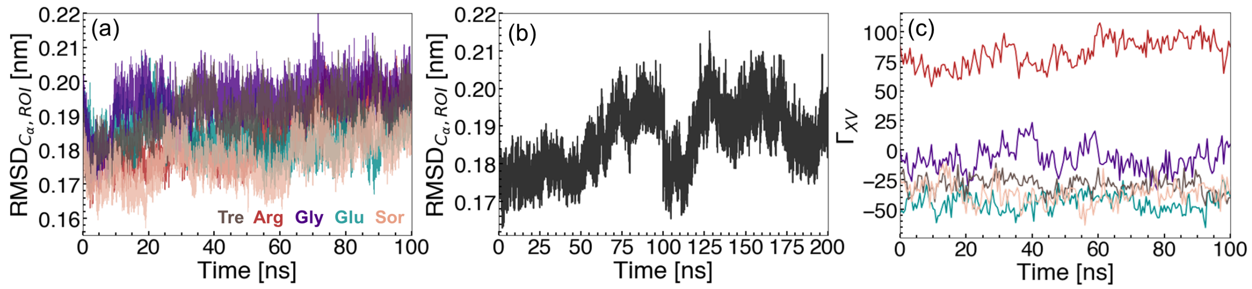


**Figure S5:** Convergence measurements for excipient simulations. (a) $C_\alpha$-RMSD of ROI capsid proteins. (b) $C_\alpha$-RMSD of ROI capsid proteins for an extended (200 ns) simulation of 0.1 M Arg / 5-fold PPV surface model. (c) Preferential interaction coefficient values ($R_{cut} = 0.7$ nm) as a function of simulation time. Data are colored according to excipient identity (Gray, Tre; Red, Arg; Purple, Gly; Cyan, Glu; Salmon, Sor).

### 3 Water Structure and Dynamics

Selecting a box height, $h_{box}$, is a non-trivial task in preparing our capsid surface simulations. Too small an $h_{box}$ and the wall placed at $z = h_{box}$ will interact with the hydration shell of the capsid. Too large an $h_{box}$ and significant computational cost is incurred. To this end, we computed bulk properties of water density ($\rho$), three-body angle distributions ($\theta$), and reorientation correlation functions ($C(t)$) with varying $h_{box}$ values.

#### 3.1 Three-Body Angle Distributions

Three-body angle distributions were computed according to Monroe and Shell[S4] as the angle between two vectors connecting a central water oxygen atom to oxygens on two of its neighbors. Oxygens are considered neighboring if they are within a 0.34 nm cut-off distance from the central water oxygen. To save on computational time, we considered only water molecules with a 1 nm thick slab between $r$ and $r + 1$, where $r$ is some distance from the capsid atom closest to $h_{box}$. Specifically, $\theta$ is obtained as:

$$\theta = \cos^{-1}\left(\frac{r_{1,2} \cdot r_{1,3}}{||r_{1,2}|| * ||r_{1,3}||}\right) \tag{S9}$$

#### 3.2 Water Reorientation Dynamics

The reorientational dynamics of water was measured by computing correlation function, $C(t)$, of the water dipole vector, $\mu$:[S5–S7]

$$C(t) = \frac{\langle \mu_i(0) \cdot \mu_i(t) \rangle}{\langle \mu_i(0) \cdot \mu_i(0) \rangle} \tag{S10}$$

where $\mu_i(t)$ is the dipole vector of the $i$th water molecule at time $t$. Water molecules were considered for analysis according to the following criteria:[?] (i) the Cartesian coordinates of the water oxygen is within $r$ to $r + dr$ at $t = 0$, (ii) if a water molecule moves into a buffer region, spanning $r + dr$ to $r + dr + b$, its position is flagged and tracked over time, and (iii) a water molecule is removed from consideration if the tracked molecule exits the buffer region without re-entry into the $r$ to $r + dr$ shell, or persists within the buffer region for at least 2 ps. In the above criteria, $r$ denotes the minimum distance from the center of the nearest capsid atom, $dr$ is the width of the hydration layer under consideration, and $b$ is the width of the buffer region. In the ensuing analyses, we use $dr = 0.5$ nm and $b = 0.1$ nm.

Additional MD simulations were performed for water dynamics analyses. Representative configurations from the end of production runs were used as initial coordinates and velocities were regenerated. These simulations were performed for 300 ps in the NVT ensemble and configurations were stored every 0.1 ps.

#### 3.3 Justification of $h_{box}$

Our criteria for a sufficiently large $h_{box}$ is that (i) images between periodic capsid images should be minimal, and (ii) a region of water should exist far enough from the wall that bulk properties are observed. We handle (i) by selecting an $h_{box}$ value at least 1.5 nm greater than the height of the capsid surface, ensuring periodic capsid surfaces are at a distance greater than the CHARMM nonbonded cut-off distance of 1.0-1.2 nm. For (ii), the decision is less straightforward. We began by computing $C(t)$ as a function of distance from the upper wall, and capsid surface, using variable $h_{box}$ values (Fig. S6). We observe that by a distance of 0.6 nm from both the capsid surface and the upper wall, C(t) stops changing as a function of distance and began to resemble bulk-like properties. With this in mind, we decided that a suitable $h_{box}$ value should be given as:

$$h_{box} = h_{capsid} + FF_{nb,cut} + 2d_{c(t)} \tag{S11}$$

where $h_{capsid}$ is the height of the capsid surface, $FF_{nb,cut}$ is the force field dependent nonbond cut-off distance, and $d_{c(t)}$ is the distance at which water dynamics are no longer perturbed by the upper wall or capsid surface. For the PPV 5-fold surface model, this worked out to

$h_{box}$ ~ 11.7 nm. For simplicity, we used a box height of 12.0 nm. Additionally, because the 5-fold surface is the largest among the PPV surface models, we applied this same 12.0 nm box height to the 2- and 3-fold surface models as well. With $h_{box}$ set to 12.0 nm, we computed $\theta$, $\rho$, and $C(t)$ at a 1 nm distance from the upper wall (Fig. S7). Here we show that in these systems, bulk properties of TIP3P water are recovered, indicating a region that is non-perturbed by either the upper wall or the capsid surface.

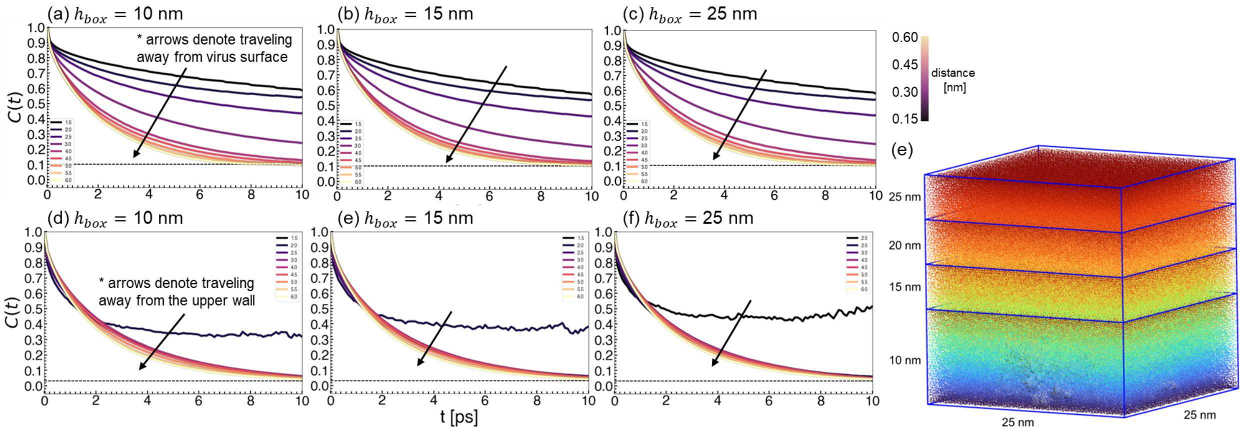


**Figure S6:** Water reorientation correlation functions for 5-fold capsid surfaces. Top row: Comparison of changing $z$-coordinates as a function of distance from the capsid surface. Bottom row: Comparison of changing $z$-coordinates as a function of distance from the upper wall. Systems with variable $h_{box}$ values of (a,d) 10 nm, (b,e) 15 nm, and (c,f) 25 nm were studied. (e) Relative heights of different $h_{box}$ values.

We additionally implemented the Yeh-Berkowitz correction[S8] without a vacuum layer, as no significant differences in water dipole distributions were observed in the presence or absence of the vacuum (Fig. S8). Because the GROMACS domain decomposition algorithm occurs even over vacuum space, our simulation speed dropped from ~51.6 ns/day to ~13.4 ns/day with a vacuum layer in the z-dimension. Because the dipole orientations of water were not significantly perturbed in the absence of the vacuum layer, we opted for the more efficient simulation configuration.

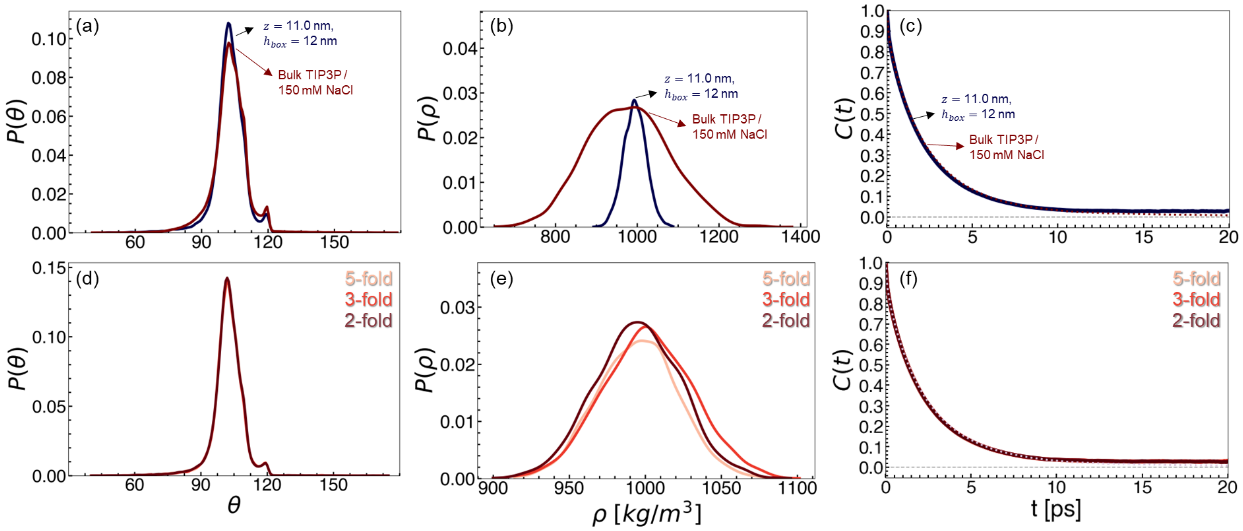


**Figure S7:** Water structure and dynamics validations for capsid systems. Top row: Comparison between water 1 nm away from the upper wall and bulk TIP3P water. Bottom row: Comparison across PPV surfaces under study (5-fold, 3-fold, and 2-fold). Data are shown for (a,d) three-body angle distributions, (b,e) density distributions, and (c,f) water dipole vector correlation functions.

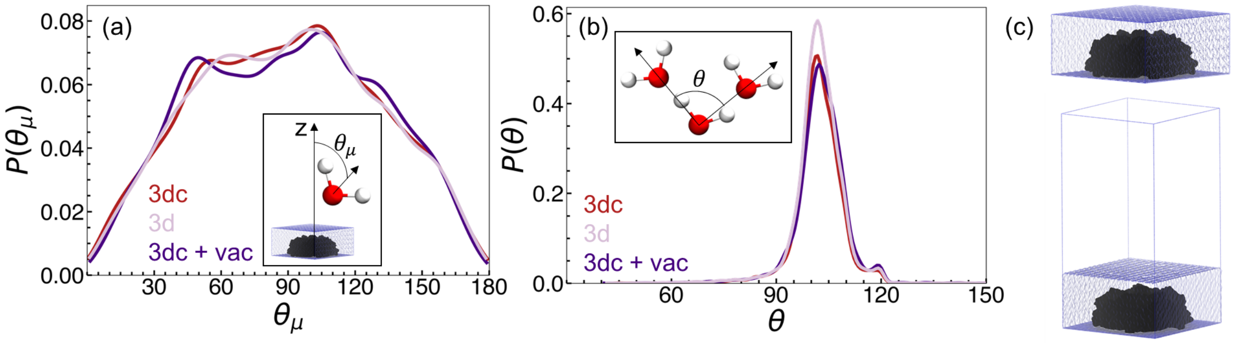


**Figure S8:** Water dipole distributions and three-body angle distributions. (a) Angular distributions are shown for water dipole vectors relative to the z-axis. (b) Three-body angle distributions for water molecules. (c) Representative snapshots of systems without (top) and with (bottom) an additional vacuum layer in the z-direction. Data are shown for systems with the Yeh-Berkowitz correction but no vacuum (red; 3dc), without the correction and with no vacuum (pink; 3d), and with both the correction and vacuum layer (purple; 3dc + vac).

### 4 Pulling Simulations

We assessed the sensitivity of capsid surface pulling simulations to parameters of box size, pulling rate, and force constant (Fig. S9). Parameters were tested on the PPV 5-fold surface model. We ultimately found that a pulling rate of 0.001 nm ps$^{-1}$ and a force constant of 1000 kJ mol$^{-1}$ nm$^2$ gave rise to a sharp decline in the fraction of native interfacial contacts after ∼1.5 ns. At slower pulling rates or smaller force constants, we did not observe clear crack onsets and capsid fragmentation. With faster pulling rates and larger force constants, the capsid broke too quickly to sufficiently sample fragmentation at the residue level. With these parameters, increasing the box size in either the $xy$ or $z$ dimensions did not qualitatively change the fragmentation behavior of the capsid surface. For box size effects (Fig. S9a), pulling rate and force constant were fixed at 0.001 nm ps$^{-1}$ and 1000 kJ mol$^{-1}$ nm$^2$, respectively. For pulling rate effects (Fig. S9b), box size was fixed at 25x25x12 nm and force constant was fixed at 1000 kJ mol$^{-1}$ nm$^2$. For force constant effects (Fig. S9c), box size was fixed at 25x25x12 nm and pulling rate was fixed at 0.001 nm ps$^{-1}$.

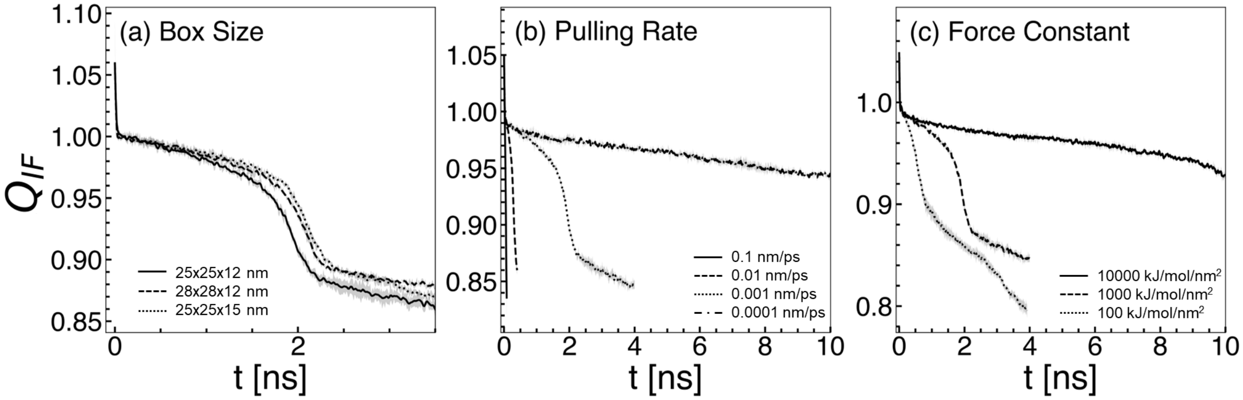


**Figure S9:** Sensitivity of the PPV 5-fold surface to different pulling parameters. The effects of (a) box size, (b) pulling rate, and (c) force constant were explored.

Within the first 10 ps of all pulling simulations, a small amount of transient interfacial contacts are broken due to random fluctuations. Hence, we shift $Q_{IF}$ curves to the left such that $Q_{IF}(\tau) = 1.0$. To capture only the response of the capsid surfaces to the applied pulling forces, analyses are carried out over the portion of the curve for which $Q_{IF} \leq 1$.

The choice of initial configurations also influences the correlation with experimental stability data (Fig. S10). As we were interested in developing a model that could accurately reproduce relative excipient effects on PPV, we decided on a pulling protocol with the strongest correlation with experimental data. In this case, it was the condition where pulling simulations were run at 300 K, pulling forces were applied to all capsid proteins, and intiial configurations were extracted after equilibrium, unrestrained molecular dynamics simulations (Fig. S10d).

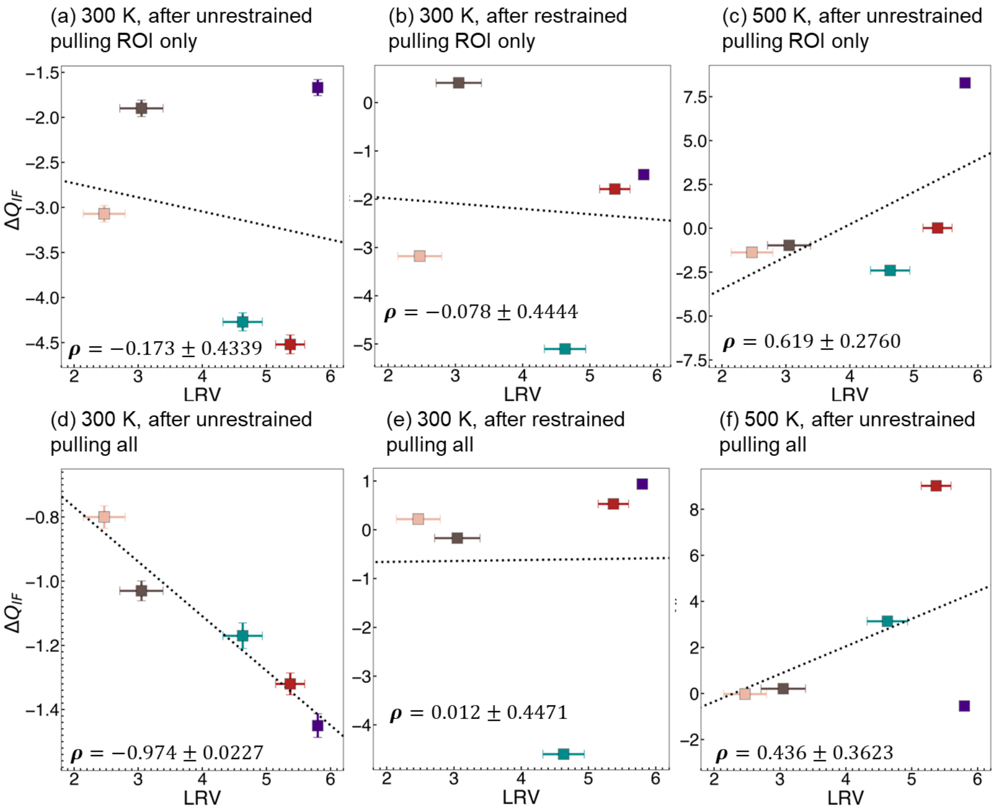


**Figure S10:** Sensitivity of the correlation of nanofragmentation results to experimental results. Conditions tested were (a) initial configurations extracted after unrestrained simulations; 300 K; pulling forces applied to only ROI proteins, (b) after restrained simulations; 300 K; pulling forces applied to only ROI proteins, (c) after unrestrained simulations; 500 K; pulling forces applied to ROI proteins only, (d) after unrestrained simulations; 300 K; pulling forces applied to all proteins, (e) after restrained simulations; 300 K; pulling forces applied to all proteins, and (f) after unrestrained simulations; 500 K; pulling forces applied to all proteins. Pearson correlation coefficients are provided in the lower left of each plot.

With a slower pulling velocity (0.0001 nm $ps^{-1}$) results in the same rank-ordering of capsid interfacial stability (e.g., 5-fold > 3-fold > 2-fold), as shown in Fig. S11a. Excipient behavior is different, however, with a slower pulling velocity (Fig. S11b), and the correlation with experimental results is reduced (Fig. S11c).

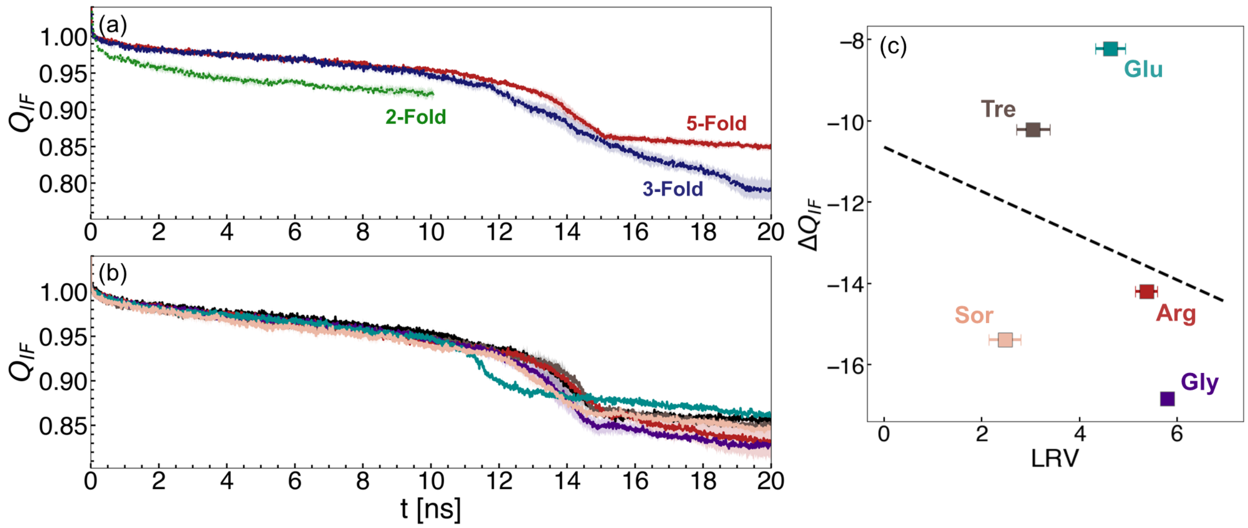


**Figure S11:** Pulling simulations with a pulling velocity of 0.0001 nm/ps. (a) Change in $Q_{IF}$ for different surface models in 0.15 M NaCl. Colors indicate 5-fold, red; 3-fold, blue; and 2-fold, green. (b) Change in $Q_{IF}$ for the PPV 5-fold surface in 0.1 M sorbitol (SOR; pink), 0.1 M trehalose (TRE; brown), 0.1 M glutamate (GLU; cyan), 0.1 M arginine (ARG; red), and 0.1 M glycine (GLY; purple). (c) Correlation between the sum of $Q_{IF}^{Ref} - Q_{IF}^{Exc}$, reported as $\Delta Q_{IF}$, and LRV for excipient solutions.

To ensure compatibility between non-equilibrium pulling simulations and the rate of solvent diffusion, we computed the average number of water molecules within 0.4 nm of an interfacial contact pair during pulling simulations. Fig. S12 shows the change in water around these contacts against the change in fraction of native contacts. These data demonstrate that hydration and capsid fragmentation occur cooperatively, and no transient cavities form during these simulations. Further, this cooperativity does not depend on the pulling rate, with similar trends observed using a pulling rate of 0.0001 nm $ps^{-1}$ (Fig. S13).

$Q_{IF}$ is observed to steadily decrease during NVT production runs (Fig. S14). The rate of decrease is consistent with the decrease in $Q_{IF}$ during the initial phase of the pulling simulations. This indicates that the combination of pulling rate and force may not significantly alter the native fluctuations in $Q_{IF}$ until the maximum force is reached, after which the crack is formed. Native interfacial contacts are observed to drop by ∼10% during equilibrium simulations. Thus, drops of greater than 10% observed in pulling simulations may be considered significant.

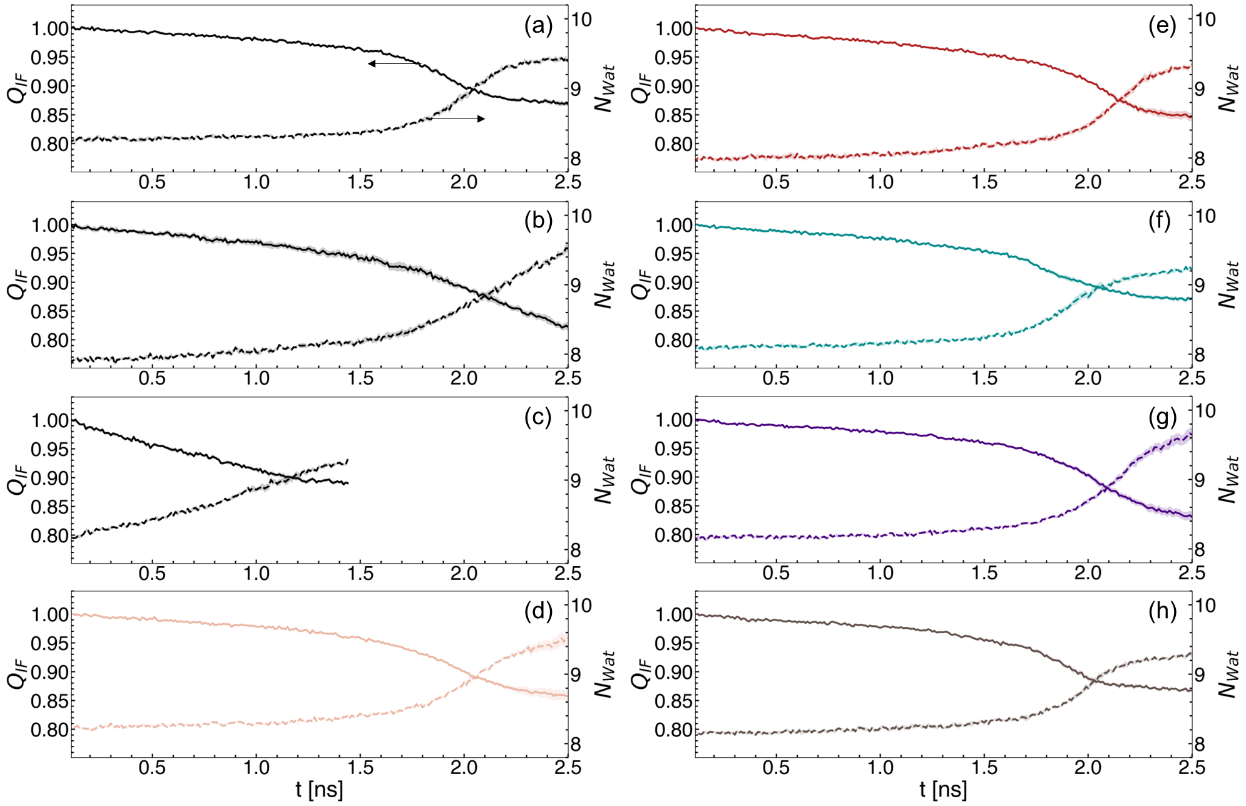


**Figure S12:** Change in the fraction of native interfacial contacts, $Q_{IF}$, left y-axes; and change in hydration waters near native interfacial contacts, $N_{Wat}$, right y-axes, with simulation time. Data are shown for pulling velocities of 0.001 nm ps$^{-1}$. In panel (a), arrows are drawn to indicate which y-axis the data belong to. (a) PPV 5-fold, (b) 3-fold, and (c) 2-fold surfaces. Data from 5-fold surfaces in excipient solutions are also shown for (d) glycerol, GOL; (e) arginine, ARG; (f) glutamate, GLU; (g) glycine, GLY; and (h) trehalose, TRE.

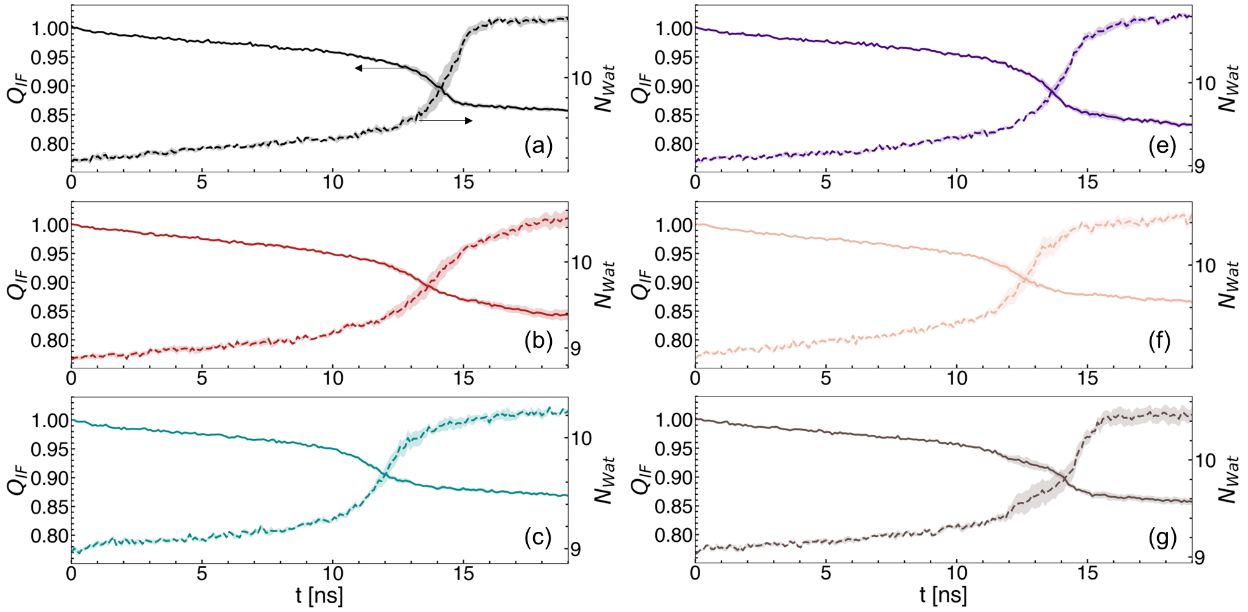


**Figure S13:** Change in the fraction of native interfacial contacts, $Q_{IF}$, left y-axes; and change in hydration waters near native interfacial contacts, $N_{Wat}$, right y-axes, with simulation time. Data are shown for pulling velocities of 0.0001 nm ps$^{-}$1. In panel (a), arrows are drawn to indicate which y-axis the data belong to. (a) PPV 5-fold surface. Data from 5-fold surfaces in excipient solutions are also shown for (b) arginine, ARG; (c) glutamate, GLU; (d) glycine, GLY; (f) glycerol, GOL; and (g) trehalose, TRE.

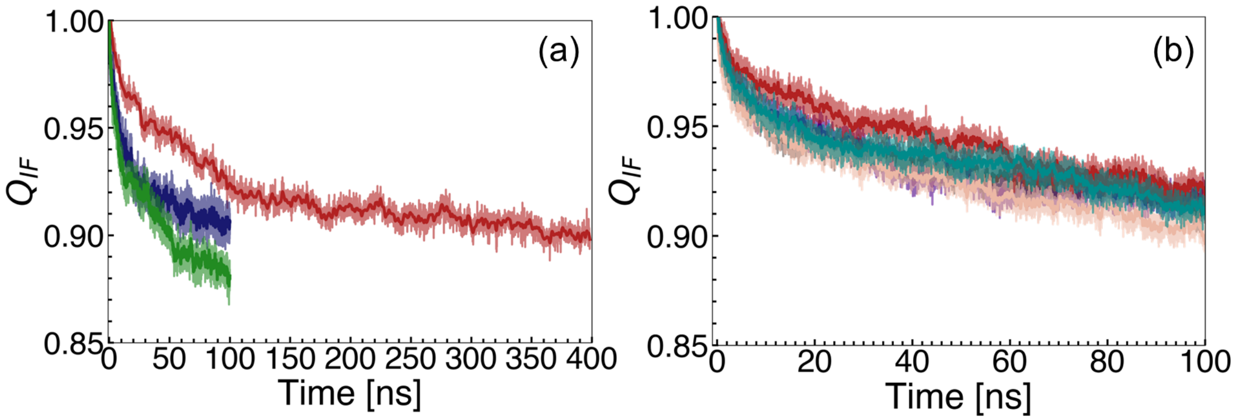


**Figure S14:** Change in the fraction of native interfacial contacts, $Q_{IF}$, with simulation time during equilibrium NVT simulations. (a) Data from 5-fold (red), 3-fold (blue), and 2-fold (green) surfaces. (b) Change in $Q_{IF}$ for the PPV 5-fold surface in 0.1 M sorbitol (SOR; pink), 0.1 M trehalose (TRE; brown), 0.1 M glutamate (GLU; cyan), 0.1 M arginine (ARG; red), and 0.1 M glycine (GLY; purple).

### 5 Performance

Performance of the assembled capsid compared to surface models is shown in Fig. S15. The performance of various capsid surface models on limited computational resources are provided in Table S2.

**Table S2:** Simulation performance and computational cost associated with various surface models tested. For all systems where performance is reported, computational resources include 24 cores and 2 v100 GPUs.

| PDB | Slice [%] | Size [atoms] | x,y [nm] | z [nm] | Performance [ns/day] |
|---|---|---|---|---|---|
| 1k3v (PPV) | 70 | 745,274 | 25.066 | 12.000 | 6.614 |
| 3j31 (STIV) | 78 | 1,385,564 | 24.367 | 24.271 | 5.215 |
| 2gsy (IBV) | 40 | 1,544,390 | 29.039 | 18.299 | 4.531 |
| 3jb8 (MCMV) | 65 | 1,389,126 | 31.210 | 14.177 | 4.906 |
| 3j6r (HPV) | 82 | 1,432,766 | 33.802 | 13.429 | 5.582 |
| 7qwz (SCV) | 74 | 1,634,369 | 35.033 | 13.681 | 4.410 |

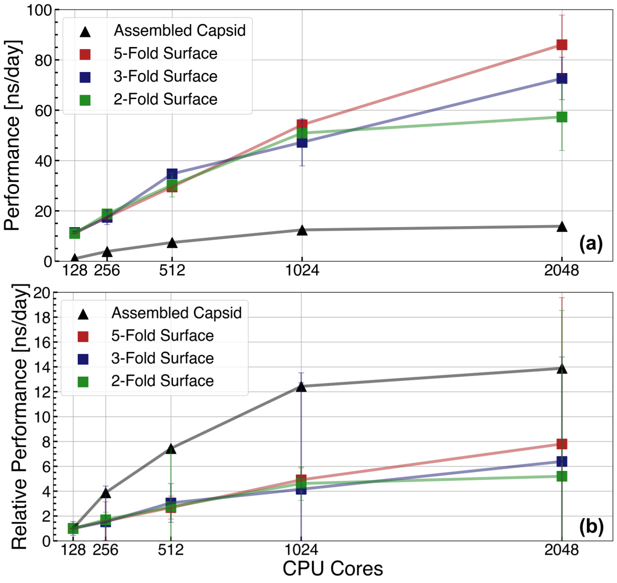


**Figure S15:** PPV simulation performance as a function of the number of available CPU cores. (a) Raw performance reported in ns/day. (b) Relative performance compared to the 128 core configuration. Colors indicate the 5-fold (red), 3-fold (blue), and 2-fold (green) surface models (squares), as well as the fully assembled PPV capsid (black triangles).

### 6 Capsid Structure and Dynamics

The 3-fold (Fig. S16f) and 2-fold (Fig. S16i) capsid surface models closely resemble the residue-residue correlations observed in the respective assembled capsid ROIs. Peripheral proteins (Fig. S16c,g,k) and restrained proteins (Fig. S16d,h,l), on the other hand, show altered residue-residue correlations. This highlights the need for setting the proper molecular context of capsid proteins.

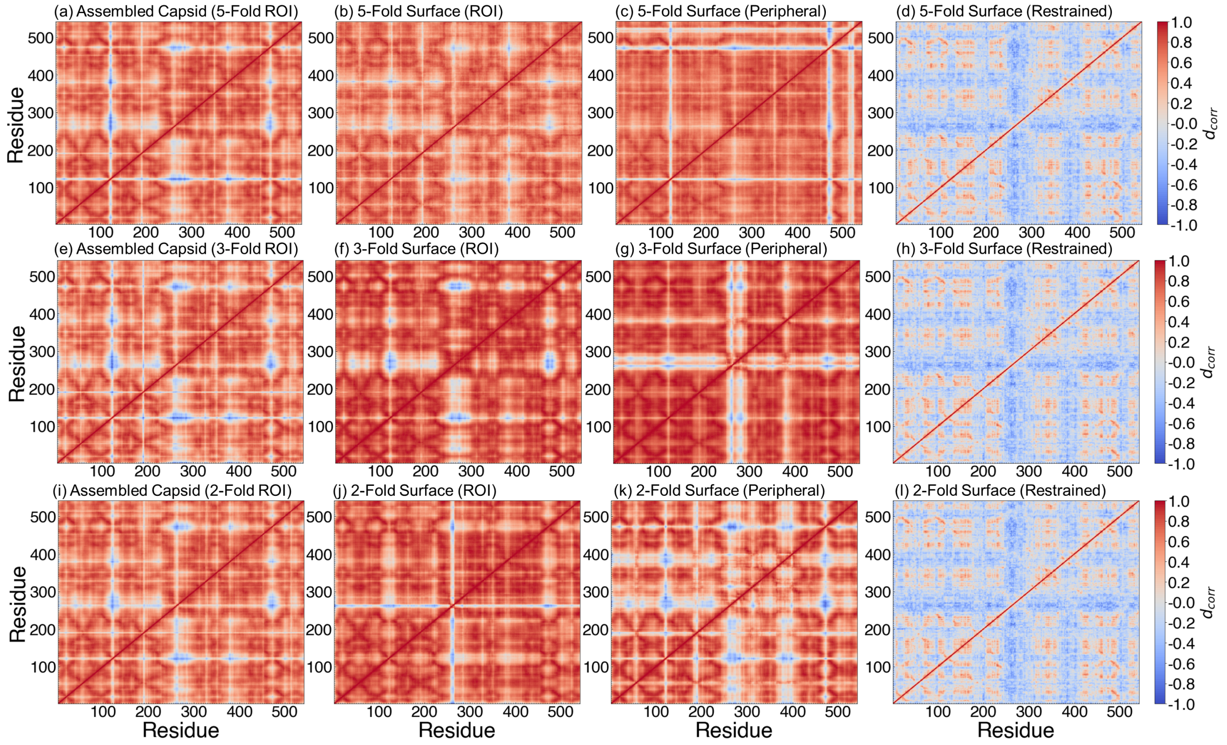


**Figure S16:** Dynamic cross-correlation matrices of residue-residue pairs within PPV capsid proteins. Top: 5-fold capsid proteins taken from the production runs of (a) the Assembled Capsid, (b) the ROI of the 5-fold Surface model, (c) the peripheral portion of the 5-fold Surface model, and (d) the restrained 5-fold Surface model. Middle: 3-fold capsid proteins taken from the production runs of (e) the Assembled Capsid, (f) the ROI of the 3-fold Surface model, (g) the peripheral portion of the 3-fold Surface model, and (h) the restrained 3-fold Surface model. Bottom: 2-fold capsid proteins taken from the production runs of (i) the Assembled Capsid, (j) the ROI of the 2-fold Surface model, (k) the peripheral portion of the 2-fold Surface model, and (l) the restrained 2-fold Surface model. $d_{corr}$ values are colored from blue to white to red, with blue indicating strong negative correlation, red indicating strong positive correlation, and white indicating weak or no correlation.

In addition to characterizing intraprotein dynamics, we assess intraprotein structure by constructing residue interaction networks (RINs) for the PPV systems under study (Fig. S18). RINs provide graph-based representations of pairwise residue-residue interaction networks within proteins and have been useful in providing insights into factors that drive protein structure, function, and stability.[S9–S12] Within each RIN, we compute betweenness centrality, $C_b$, as a measure for how often a given residue acts as a bridge connecting all other residues. Several key residues with high $C_b$ values are identified (Fig. 5a-c): Tyr185, Pro358, Asn370, Arg404, Thr456, and Gln493, indicating significance of these residues in forming the core of the PPV capsid protein. Importantly, these key residues are identified in both the assembled PPV capsid and 5-fold surface model. The networks of residue-residue interactions are similar between the ROIs from the assembled capsid and capsid surface (Fig. 5d). Persistent interactions are made among protrusion-forming residues, as well as between protrusion- and dimple-forming residues. Together, these data show that the capsid surface models generated through our workflow accurately capture the same network of intraprotein interactions as in the assembled PPV capsid, regardless of the ROI.

We report distributions of sphericity, $\Psi$ as a global measure of structure. $\Psi$ measures the deviation of each capsid model from well-characterized shapes, such as hemispheres,

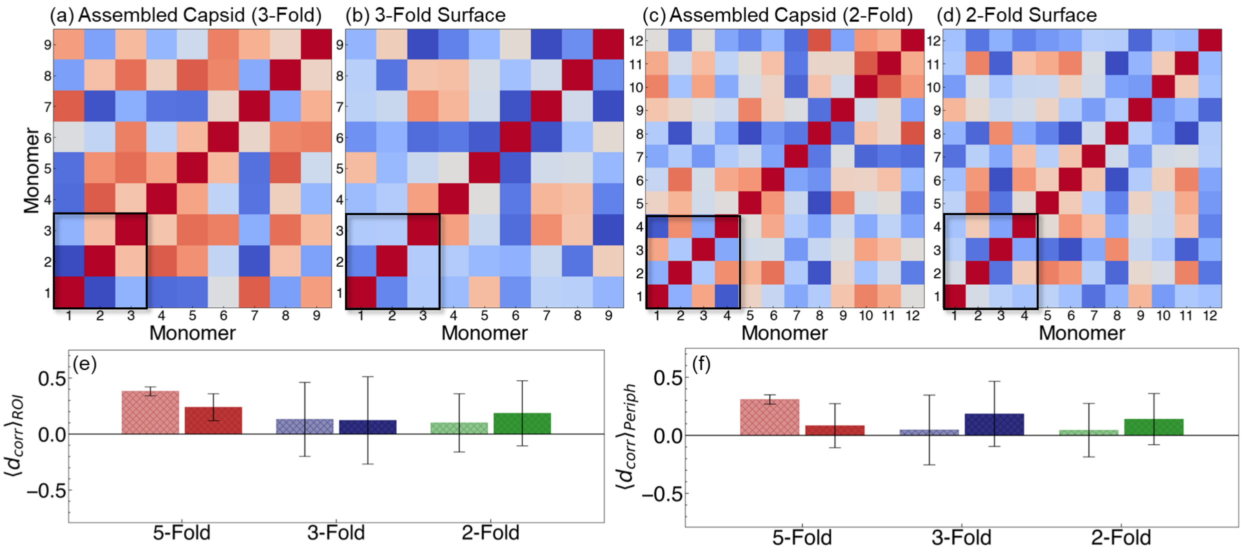


**Figure S17:** Dynamic cross-correlation matrices of monomer-monomer pairs within PPV capsids. (a) 3-fold capsid proteins from the assembled capsid ROI, (b) 3-fold proteins from the 3-fold surface model, (c) 2-fold proteins from the assembled capisd ROI, (d) 2-fold proteins from the 2-fold surface model. In (e) and (f), the mean and standard deviation for either (e) ROI or (f) peripheral capsid proteins are reported for each system.

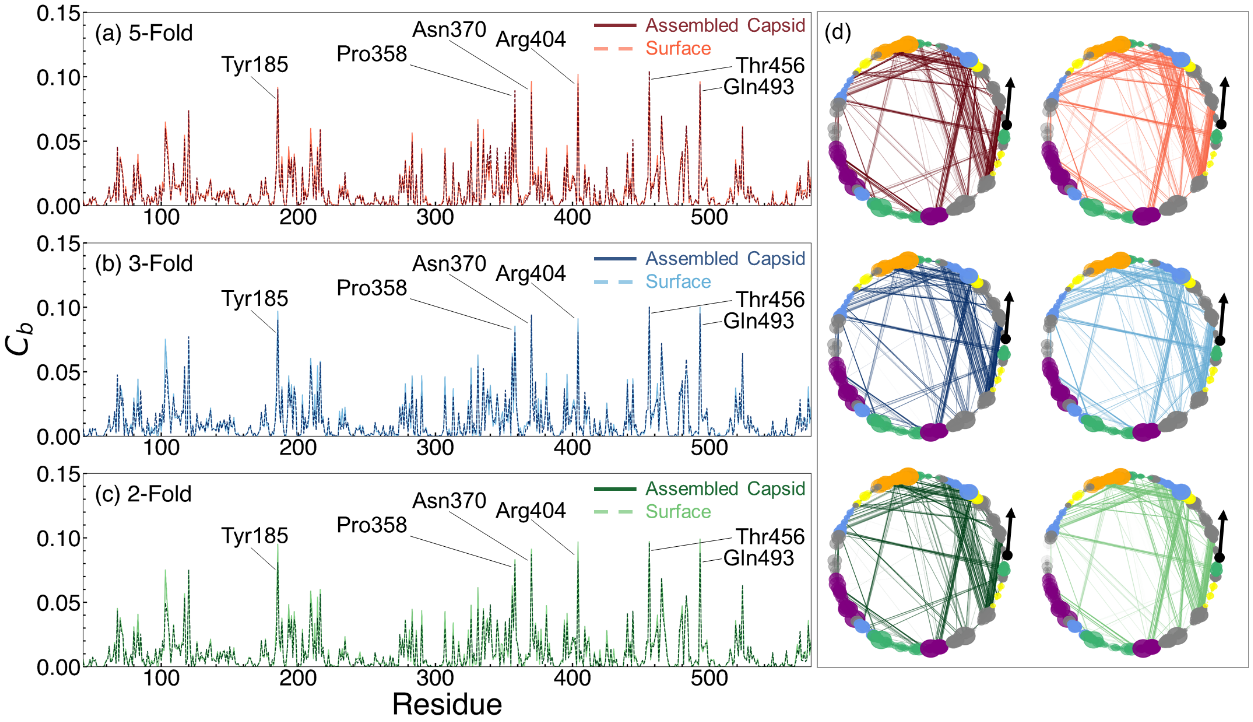


**Figure S18:** Residue interaction network analysis of PPV systems under study. Betweenness centrality ($C_b$) values are plotted for residues at the (a) 5-fold, (b) 3-fold, and (c) 2-fold ROIs. ROIs from the assembled capsid are shown as dark solid lines, while capsid surface ROIs are shown as light dashed lines. (d) Graph representation of intraprotein residue interaction networks. Nodes represent residues and are colored according to surface feature (buried, gray; 5-fold pore, orange; 5-fold canyon, yellow; 3-fold exterior protrusion, blue; 3-fold interior protrusion, purple; 2-fold dimple, green). Edges are drawn between nodes where residues are in contact for >80% of the simulation time. Edge widths are proportional to the total amount of time two residues are in contact ($r_{ij} < 0.4$ nm). Node sizes are proportional to residue betweenness centrality. The N-terminus is marked with an arrow that shows the direction of the amino acid sequence.

icosahedrons, and spheres. The fully assembled capsid has a $P(\Psi)$ distribution centered somewhere between an icosahedron and a perfect sphere (Fig. S19). This sphericity value for an icosahedral capsid shell is similar to that of HBV, obtained by Hadden *et al.*[S13]

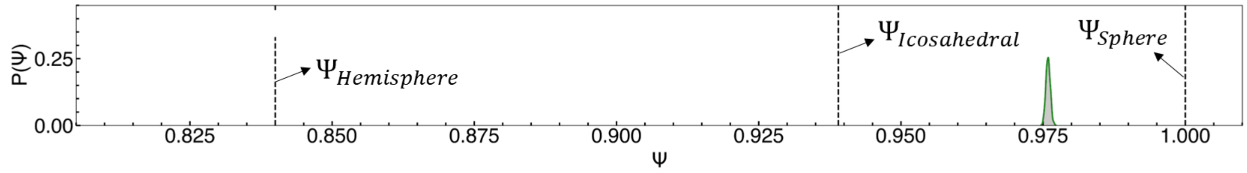


**Figure S19:** Sphericity distribution of the fully assembled capsid. Vertical dashed lines denote typical $\Psi$ values for hemispheres, icosahedrons, and spheres, from left to right.

### 7 Experimental Details

##### Materials

Eagle's minimum essential media (EMEM), sodium bicarbonate (7.5% solution), penicillin/streptomycin (pen/strep, 10,000 U/ml), fetal bovine serum (FBS, qualified, USDA-approved regions), phosphate-buffered saline (1 X PBS, pH 7.2), and trypsin/EDTA (0.25%) used for cell culture were purchased from Gibco™ (Grand Island, NY). MTT (2-(3,5-diphenyltetrazol-2-ium-2-yl)- 4,5-dimethyl-1,3-thiazole; bromide, 98%) and sodium dodecyl sulfate (SDS, BioReagent, ≥98.5%) were purchased from Fisher Scientific (Waltham, MA) for virus titration. Arginine monohydrochloride (reagent grade, ≥98% (HPLC)) was purchased from Millipore Sigma (Burlington, MA). D-(+)-Trehalose dihydrate from Saccharomyces cerevisiae (>99%) was purchased from Sigma-Aldrich (St. Louis, MO). L-glutamic acid monosodium salt (cell culture reagent) was purchased from MP Biomedicals (Santa Ana, CA). Glycine (>95%) was purchased from Fisher Scientific (Hampton, NH). D-sorbitol (100%) was were purchased from Thermo Fisher Scientific Chemicals (Waltham, MA) Sodium phosphate monobasic monohydrate (reagent ACS grade) was purchased from Millipore. Sodium phosphate dibasic heptahydrate (ACS reagent, ≥98.0%) was purchased from Sigma Aldrich (St. Louis, MO).

##### Methods

*Cell line and virus*

Porcine kidney cells (PK-13) were purchased from the American Type Culture Collection (ATCC®) (cat# CRL-6489™) and cultured in EMEM supplemented with 10 v/v% FBS and 1 v/v% pen/strep. The cells were incubated at 37 °C, 5% $CO_2$, and 100% relative humidity. Porcine parvovirus (PPV) strain NADL-2 was a generous gift from Dr. Ruben Carbonell at North Carolina State University (Raleigh, NC). PPV strain NADL-2 was propagated in PK-13 cells using a previously established method.[S14] After three freeze-thaw cycles from -20 °C to room temperature, the cells were scraped and the supernatant was clarified by centrifugation at 5,000 rpm at 4 °C for 15 minutes in an ST16R centrifuge (Thermo Scientific

(Waltham, Ma)) with a TX-400 swing-bucket rotor. The PPV-containing supernatant was stored at -80 °C prior to use.

*Virus quantification*

The titer of PPV was found by the MTT colorimetric cell viability assay.[S15] PK-13 cells were seeded at a density of $8 \times 10^4$ cells/mL in 96-well plates and incubated overnight. The next day, the cells were infected with a 1:5 serial dilution of samples. After six days, 5 mg/mL of MTT in 1X pH 7.2 PBS was added to each well. Four hours later, 10 w/v% SDS with 0.01 M hydrochloric acid (HCl) was added to each well and the absorbance at 550 nm was measured the next day on a Synergy$^{TM}$ Mx microplate reader from BioTek (Winoski, VT). The 50% viral infectious dose was determined in units of $MTT_{50}$/mL.

*Liquid viral sample preparation*

The excipient solutions were made by dissolving different concentrations of 0.7 M arginine monohydrochloride, monosodium glutamate, glycine, trehalose, or sorbitol in phosphate buffer containing 1.54 mM sodium phosphate monobasic monohydrate and 2.71 mM sodium phosphate dibasic (pH 7.2). The virus samples were made by adding 10 v/v% viral stock solutions to the excipient solution.

*Thermostability studies*

Liquid samples were prepared in triplicate and were put either in a heat block at 60 °C[S16] or in a fridge at 4 °C as the control samples. 72 hours later, the titer of virus in each sample was determined using the MTT assay.

### References

(S1) Inoue, H.; Timasheff, S. N. Preferential and absolute interactions of solvent components with proteins in mixed solvent systems. *Biopolymers* **1972**, *11*, 737–743.

(S2) Record, M.; Anderson, C. Interpretation of preferential interaction coefficients of nonelectrolytes and of electrolyte ions in terms of a two-domain model. *Biophysical Journal* **1995**, *68*, 786–794.

(S3) Shukla, D.; Shinde, C.; Trout, B. L. Molecular Computations of Preferential Interaction Coefficients of Proteins. *J. Phys. Chem. B* **2009**, *113*, 12546–12554.

(S4) Monroe, J. I.; Shell, M. S. Decoding signatures of structure, bulk thermodynamics, and solvation in three-body angle distributions of rigid water models. *The Journal of Chemical Physics* **2019**, *151*, 094501.

(S5) Liu, H.; Xiang, S.; Zhu, H.; Li, L. The Structural and Dynamical Properties of the Hydration of SNase Based on a Molecular Dynamics Simulation. *Molecules* **2021**, *26*, 5403.

(S6) Diaz, A.; Ramakrishnan, V. Effect of osmolytes on the EcoRI endonuclease: Insights into hydration and protein dynamics from molecular dynamics simulations. *Computational Biology and Chemistry* **2023**, *105*, 107883.

(S7) Santra, S.; Jana, M. Influence of Aqueous Arginine Solution on Regulating Conformational Stability and Hydration Properties of the Secondary Structural Segments of a Protein at Elevated Temperatures: A Molecular Dynamics Study. *J. Phys. Chem. B* **2022**, *126*, 1462–1476.

(S8) Yeh, I.-C.; Berkowitz, M. L. Ewald summation for systems with slab geometry. *The Journal of Chemical Physics* **1999**, *111*, 3155–3162.

(S9) Grewal, R.; Roy, S. Modeling proteins as residue interaction networks. *PPL* **2015**, *22*, 923–933.

(S10) Franke, L.; Peter, C. Visualizing the Residue Interaction Landscape of Proteins by Temporal Network Embedding. *J. Chem. Theory Comput.* **2023**, *19*, 2985–2995.

(S11) Dasetty, S.; Zajac, J. W. P.; Sarupria, S. Exploitation of active site flexibility-low temperature activity relation for engineering broad range temperature active enzymes. *Mol. Syst. Des. Eng.* **2023**, *8*, 1355–1370.

(S12) Yehorova, D.; Di Geronimo, B.; Robinson, M.; Kasson, P. M.; Kamerlin, S. C. Using residue interaction networks to understand protein function and evolution and to engineer new proteins. *Current Opinion in Structural Biology* **2024**, *89*, 102922.

(S13) Hadden, J. A.; Perilla, J. R.; Schlicksup, C. J.; Venkatakrishnan, B.; Zlotnick, A.; Schulten, K. All-atom molecular dynamics of the HBV capsid reveals insights into biological function and cryo-EM resolution limits. *eLife* **2018**, *7*, e32478.

(S14) Heldt, C. L.; Hernandez, R.; Mudiganti, U.; Gurgel, P. V.; Brown, D. T.; Carbonell, R. G. A colorimetric assay for viral agents that produce cytopathic effects. *Journal of Virological Methods* **2006**, *135*, 56–65.

(S15) Joshi, P. U.; Decker, C.; Zeng, X.; Sathyavageeswaran, A.; Perry, S. L.; Heldt, C. L. Design Rules for the Sequestration of Viruses into Polypeptide Complex Coacervates. *Biomacromolecules* **2024**, *25*, 741–753.

(S16) Mi, X.; Blocher McTigue, W. C.; Joshi, P. U.; Bunker, M. K.; Heldt, C. L.; Perry, S. L. Thermostabilization of viruses *via* complex coacervation. *Biomater. Sci.* **2020**, *8*, 7082–7092.
